# A toolkit to identify parameter degeneracy and solution manifolds of Hodgkin-Huxley type current models

**DOI:** 10.64898/2026.11.16.767143

**Authors:** Saina Namazifard, Fabrizio Gabbiani, Matthias Heinkenschloss

## Abstract

Countless studies have fitted Hodgkin-Huxley (HH) type models of active currents to voltage clamp data using nonlinear least squares. Yet, little is known about the uniqueness of parameter solutions and the interdependence of HH-type model parameters. Here, we show that considerable insight can be gained from the singular value decomposition (SVD) of a matrix closely related to the objective function used in the fitting procedure. After fitting all model parameters simultaneously using synthetic data originating from a representative range of experimental cases, this reveals that a manifold of solutions is possible depending on the type of voltage clamp data available and on the selected HH-type model. In such cases, we use the SVD to determine which parameters are least identifiable. Next, we develop an SVD-based manifold tracing technique that can efficiently unveil the structure of the manifold of equivalent solutions in the parameter space of the model. Our methods also identify parameter interdependencies and their impact on joint parameter confidence regions. The most salient one consistently observed across our experimental examples was between the peak conductance and reversal potential of active HH-type current models. The SVD-based strategy presented here is expected to be generally applicable to investigate parameter degeneracy and solution manifold structure in biophysical models of single neurons. Thus, our results should prove valuable for understanding the constraints imposed by the biophysics of ion channels on single neuron computation.

**Author summary:** Mathematical modeling is extensively used to describe how neurons generate electrical signals. It is also used to fit models of ion channels from experimental data. However, a good fit does not mean that the estimated model parameters are unique. Although it is known that different combinations of model parameters can produce similar neuronal outputs, we develop a systematic approach to characterize such parameter combinations with a focus on voltage clamp experiments. Our toolkit determines which parameters are not well constrained by the experiments. It precisely traces the local solution manifold at low computational cost compared to exhaustive sampling methods, showing how different parameters compensate one another to generate nearly identical outputs. Estimating all parameters simultaneously also reveals parameter interdependencies and their effects on the uncertainty in their estimated values. Our results highlight the importance of understanding the limitations of voltage clamp data when estimating ion channel properties and provide a general approach for studying this issue in mathematical models of neuronal activity.

## 1 Introduction

Since the seminal work of Hodgkin and Huxley (HH), the kinetic properties of ion channels are extracted from voltage clamp experiments by fitting the parameters of first-order linear ordinary differential equations [1]. These nonlinear least squares fits are usually carried out one current trace at a time to obtain, e.g., the voltage-dependent time constant, steady-state activation, and eventual inactivation of the studied channel. While effective, this approach does not yield information on parameter dependencies across clamp potentials or on the possible non-uniqueness of the resulting model.

Parameter degeneracy – or non-uniqueness – refers to a system’s ability to produce nearly identical outputs from different parameter sets [2, 3]. While this concept has attracted little attention in the context of voltage clamp experiments, it has been extensively applied to explain the robustness of single neuron and neuronal network activity *in vivo*. Experiments on the crustacean stomatogastric ganglion (STG) – a part of the digestive nervous system – have shown that peak ionic conductances varying several folds can generate closely matching neural activity in single neurons across animals [4–8].

In compartmental models of the STG, parameter degeneracy has usually been demonstrated by exhaustive sampling across a grid of evenly spaced parameter values within their biologically feasible range [9–11] (for an alternative, see [12]). The computational cost of this approach is high and grows exponentially with the number of parameters. In addition, it provides only an approximation to the solution manifold whose resolution is determined by that of the sampling grid.

In this study, we develop methods to uncover parameter degeneracy and dependencies in HH-type current models fitted to voltage clamp data. Using variants of the Gauss-Newton method, we fit all model parameters simultaneously. After convergence, this enables us to derive the local geometry of the loss function at the solution and to assess eventual parameter degeneracy. We then determine the least identifiable parameters and develop a numerical continuation method to efficiently trace the associated manifold of degenerate parameters.

We focus on voltage clamp data for four reasons. First, identifying the parameter solution manifold of HH-type current models is an interesting problem on its own, contributing to a better understanding of how to model the underlying ion channel biophysics. Second, the properties of ion channels vary from cell to cell – and even within a cell – due to a large spectrum of potential protein variants and secondary modifications [13–16]. This makes it important to understand what can be learned from single experiments, without averaging [17]. Third, the voltage clamp assumption enables us to bypass modeling the composite voltage- and time-dependent activation dynamics of the current in a compartmental model. Since the membrane potential is held at a fixed value, its differential equation is eliminated from the compartmental model and the gating dynamics can be described analytically, simplifying the numerical determination of the solution manifold. Yet, our results show that despite this simplification, the nonlinear nature of HH-type models leads to surprising results. Fourth, our approach is expected to generalize to compartmental HH-type models with time-dependent membrane potential, provided adequate adjustments are made.

The remainder of this paper is organized as follows. In the Results, we first provide an overview of our approach, followed by the analysis of parameter degeneracy and dependencies applied to two sub-threshold currents: the H-current under two types of voltage clamp experiments, and the inactivating A-type potassium current. Next, we analyze the spike-generating fast sodium and delayed rectifier potassium current models of Hodgkin and Huxley. Following the Discussion, the Models and Methods section provides a detailed description of the techniques we used and developed. The Supporting Information provides additional graphical and mathematical results complementing the arguments of the main text. Python code implementing our results is available at Zenodo (https://doi.org/10.5281/zenodo.24285948). The DOI will become publicly accessible upon publication of the manuscript.

## 2. Results

To cover a wide range of HH-type models and experimental situations, we consider data extracted from four studies. The first one describes the hyperpolarization-activated (H-) current of the Lobula Giant Movement Detector (LGMD) neuron in grasshoppers, characterized through intracellular sharp electrode recordings [18]. The second one characterizes the H-current in rat CA1 hippocampal pyramidal cells through cell-attached patch-clamp somatic and dendritic recordings [19]. The third one studies an inactivating potassium current, the A-current, in honeybee Kenyon cells through whole-cell patch-clamp recordings [20]. The fourth one is the classic HH model for the fast sodium and delayed rectifier currents generating the action potential in the squid giant axon [1].

Each of these studies derived an HH-type model and an optimal parameter set for its studied current(s), which we used to generate synthetic voltage clamp traces that were fit as a function of the model parameters. Fitting all parameters simultaneously allows us to study their correlations. In addition, the first three studies allow us to estimate the recording noise, whereas in the HH experiments the recording noise was negligible. We used the estimated noise to assess its effect on parameter degeneracy and uncertainty.

The H-current is described by the simplest HH-type model consisting of a single activation variable, *h*. This allows for detailed mathematical results [21]. Those relevant to this study are summarized in the Supporting Information. The A-current has an activation variable, *n*, raised to the third power and an inactivation variable, *h*, so that the resulting conductance is proportional to *n*^3^*h*. The HH-model is of the functional form *m*^3^*h* for the fast sodium conductance (*m*: activation; *h*: inactivation) and *n*^4^ for the delayed rectifier (*n*: activation). In the next subsections, we show that all the experimental and modeling conditions described above impact parameter degeneracy: recording type, recording noise, and model functional form.

For the numerical non-linear least squares fitting procedure, we use variants of the Gauss-Newton method appropriate for ill-conditioned or rank-deficient problems (sec. 4.2.1). The objective function to be minimized is the sum-of-squares of the residuals, 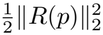, where the residual vector *R*(*p*) is the difference between the synthetic membrane currents derived from the experimental data and those generated by the model with parameters *p*. The residual *R*(*p*) has dimension *m* = *n*_*V*_ *m*_*V*_ determined by the number of fitted membrane current traces (*n*_*V*_) times the number of sampled time points per current trace (*m*_*V*_). Its derivative with respect to the *n* model parameters is the *m × n* Jacobian matrix, *J*(*p*) (with *m* ≫ *n*). The rank, *r*, of *J*(*p*) is determined by the number of non-zero singular values in its singular value decomposition (SVD, sec. 4.2.2) [22]. When *r < n*, the solution to the nonlinear least squares problem is not unique [22]. We then apply a subset selection algorithm to identify which *r* columns of the Jacobian optimally span its column space (sec. 4.2.3). The remaining *n* − *r* columns specify the least identifiable parameters.

The Jacobian can also be used to compute a first-order estimate for the matrix of second derivatives of the objective function, or Hessian. At the solution points, this procedure may be used to determine the uncertainty in the parameters (under the assumption of white noise) and their correlations (sec. 4.2.5).

### 2.1 Parameter degeneracy in a hyperpolarization-activated current model

In the HH formalism, the measured current from a voltage clamp protocol is determined by the product of the maximal conductance, the gating variable(s), and the driving force, given by the difference between the membrane potential and the ion channel reversal potential. The simplest HH conductance model comprises one activation variable, as needed to fit the ubiquitous H-current. As specified in sec. 4, we use the model eq. 1 – eq. 3 derived from voltage clamp recordings of the LGMD neuron (see [18, Fig. 2-figure supplement 1.D]). The parameters generating the target synthetic data are referred to as the ‘true parameters’ (Table S1). Plugged into eq. 4, they determine the steady-state activation and time constant (Fig. 1A). Applying the voltage clamp protocol yields the model’s target outputs, referred to as the ‘measured currents’ (Fig. 1B, colored traces).

**Fig 1.**
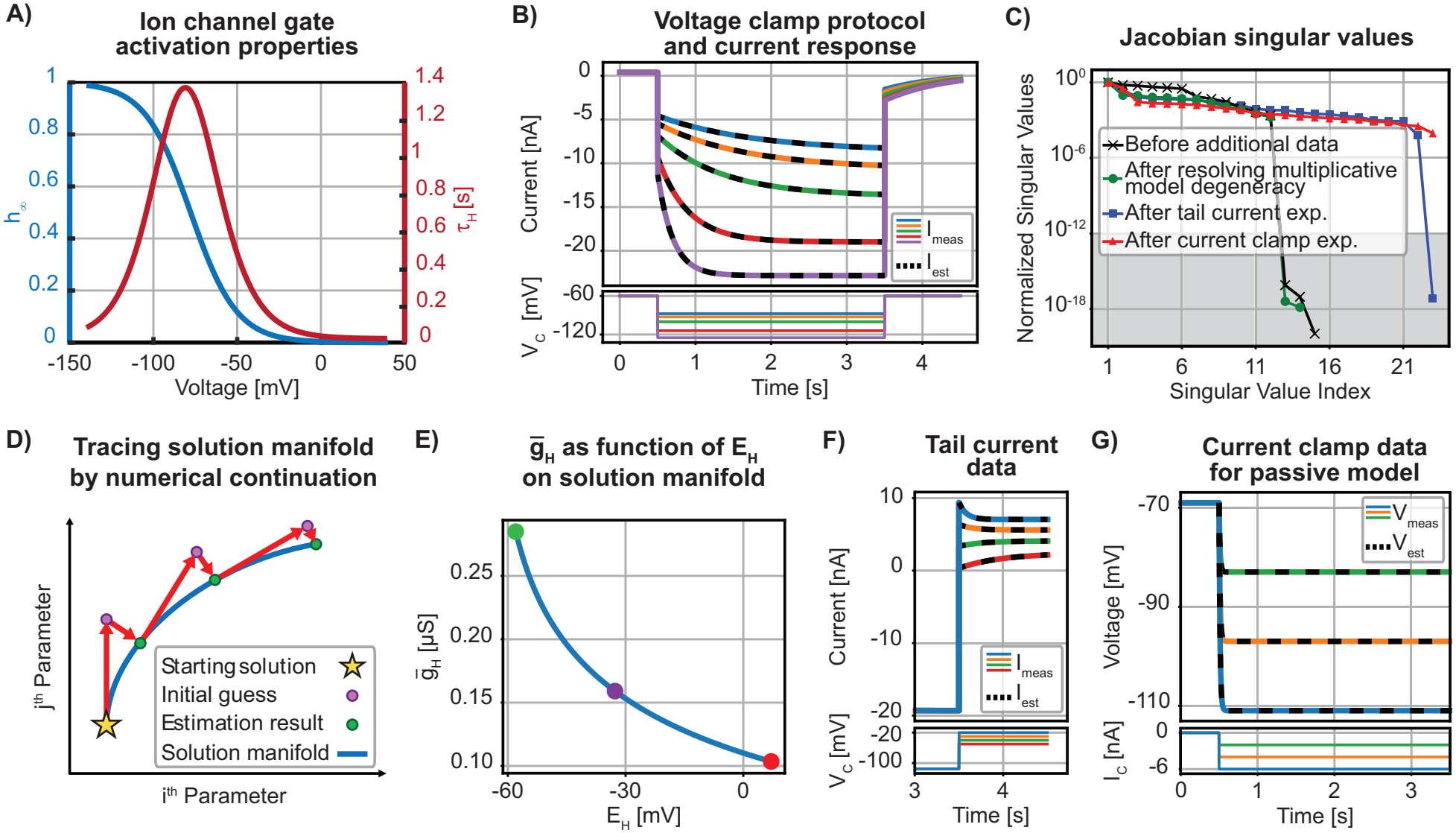
Parameter estimation and degeneracy in voltage clamp experiments for the LGMD H-current model. **(A)** Steady-state activation properties that determine the H-current dynamics. **(B)** Voltage clamp data used for parameter estimation and the corresponding model fits (colored and black dashed lines, respectively). **(C)** Normalized Jacobian singular values at different stages of the model determination process. Singular values that fall within the gray region, below the threshold of 10^−12^, are considered to be equal to zero. **(D)** Numerical continuation approach used to trace the solution manifold. **(E)** Dependence of 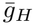 on *E*_*H*_ for fixed *E*_*L*_ on the solution manifold after resolving multiplicative model degeneracy. The three colored dots are shown to generate identical voltage clamp outputs in Fig. S1. **(F)** Tail current measurements and **(G)** current clamp recordings in a passive model (simulating pharmacological blockade of the H-current) constrain *E*_*H*_ and *E*_*L*_, respectively.

**Fig 2.**
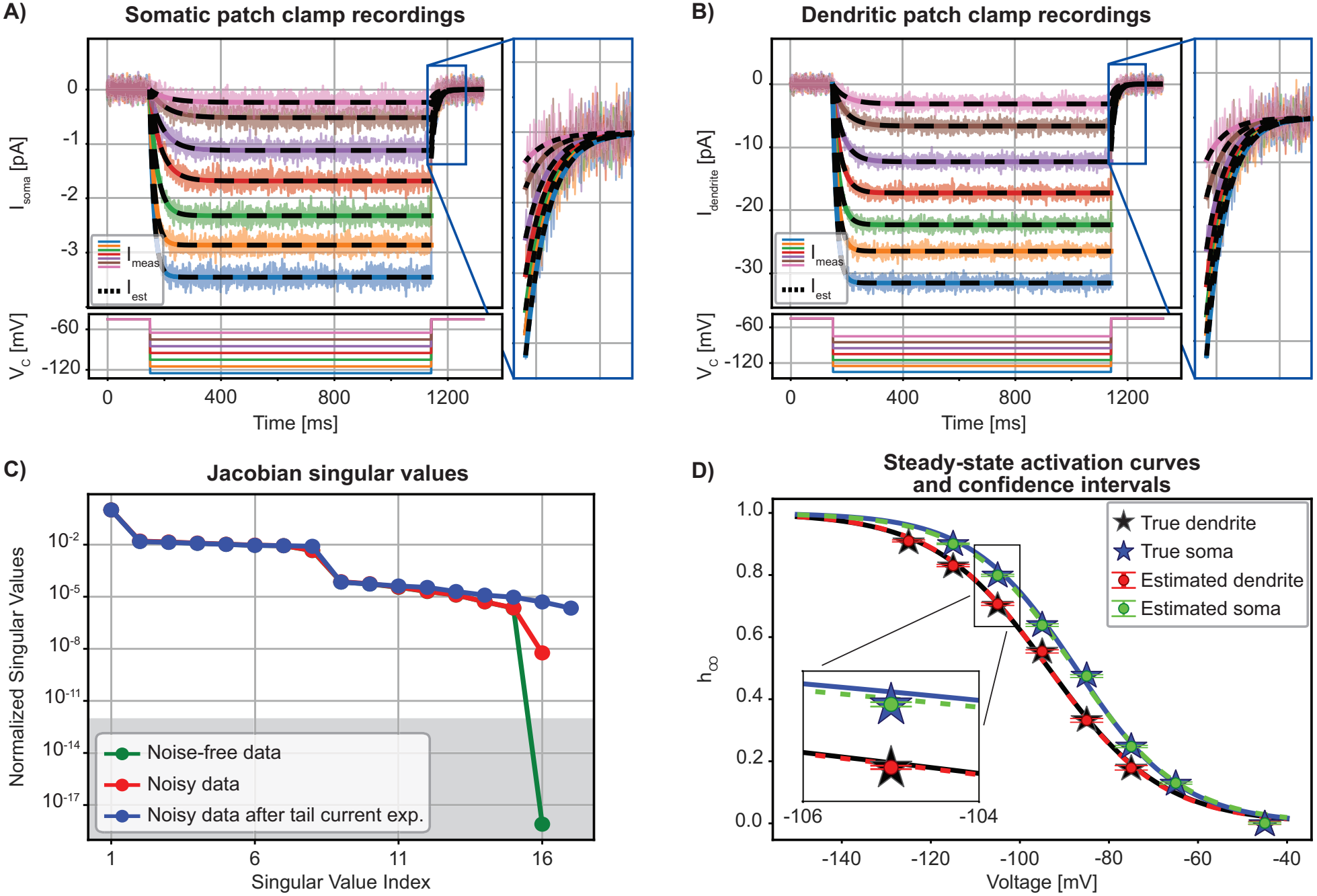
Effect of measurement noise on parameter estimation of H-current recordings from cell-attached patch-clamp experiments in CA1 pyramidal cells. **(A)** Voltage clamp recordings from the somatic membrane patch used for parameter estimation, and the corresponding model fits (colored and black dashed lines, respectively; noise represented by lighter colored shading). The tail current region is magnified on the right. **(B)** Same for the dendritic membrane patch. **(C)** Normalized Jacobian singular values for somatic recordings. Singular values that fall within the gray region, below the threshold of 10^−12^, are considered to be equal to zero. Measurement noise removes the Jacobian rank deficiency; however, the problem remains ill-conditioned with a large condition number. Adding tail current measurements to constrain *E*_*H*_ improves the Jacobian condition number. **(D)** Steady-state activation curves for soma and dendrite recordings. Activation curves are obtained by fitting a sigmoid function to the true and estimated values for each voltage clamp step. The estimated values and their corresponding 95% confidence intervals demonstrate accurate parameter estimation despite substantial measurement noise. Note that the activation values at the most negative voltage clamp step were fixed to one and excluded from the estimation; therefore, they are not shown (*h*_∞_(−125) for soma and *h*_∞_(−135) for dendrite).

We estimate the ‘unknown parameters’ by minimizing the squared norm of the residual vector from the start to the end of the voltage steps (i.e., from 0.5 to 3.5 [s] in Fig. 1B). There are fifteen parameters: the maximal conductances 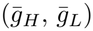, reversal potentials (*E*_*H*_, *E*_*L*_), steady-state activation at the holding potential (*h*_∞_(*V*_*C*,hold_)) and at each voltage step (*h*_∞_(*V*_*C,i*_), *i* = 1, …, 5), as well as the corresponding time constants (*τ*_*h*_(*V*_*C,i*_), *i* = 1, …, 5). However, uniqueness of the estimated parameters is not guaranteed, even after a successful fit (black dashed lines in Fig. 1B). Indeed, the Jacobian is rank-deficient by three, as its last three singular values are zero within machine precision (see sec. 4.2.2; Fig. 1C, black trace). This result can also be proven analytically and is generally valid, irrespective of the specific model parameter values (sec. S.1).

The subset selection algorithm reveals that the activation term (*h*) and the two reversal potentials (*E*_*H*_ and *E*_*L*_) are least identifiable. That *h* is least identifiable is unsurprising since, in the model formulation, it always multiplies the maximal conductance 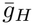, see eq. 5 in sec. 4 and [21]. As a result, only 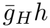 is identifiable, unless one value of *h* is fixed and excluded from the estimation problem. We set *h*_∞_(*V*_*C*,5_) = 1, assuming that the H-current is fully activated at the largest hyperpolarized voltage step (*V*_*C*,5_ = − 125 [mV]). As a result, the number of parameters is reduced to fourteen. Note, however, that if this assumption is incorrect, it will introduce a systematic error in the estimation of *h* (see sec. S.2; note that in Fig. 1A the true value is *h*_∞_(*V*_*C*,5_) = 0.968; see also sec. 2.5).

After removing this model-based multiplicative parameter degeneracy, two near-zero singular values remain, associated with *E*_*H*_ and *E*_*L*_ (Fig. 1C, green trace; see sec. S.1 for the corresponding analytical result). This is more surprising and suggests the existence of a solution manifold for the corresponding value of the objective function. To trace this manifold, we developed an iterative numerical continuation method, in which each step follows the direction in parameter space that yields the smallest change in the objective function and the resulting model output. The least-squares parameter estimation is then repeated to converge back to the solution manifold. This new parameter set serves as the starting point for the next continuation step (Fig. 1D; see sec. 4.2.4 for details). How 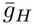 varies as a function of *E*_*H*_ along the resulting solution manifold is shown in Fig. 1E (for *E*_*L*_ = −69 [mV]). This demonstrates that a wide range of reversal potentials can be compensated by changes in other parameters, such as 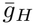, generating nearly identical voltage clamp outputs (Fig. S1; for similar results when *E*_*H*_ is fixed, see Fig. S2). Thus, to alleviate the non-uniqueness of the solution, either *E*_*H*_ and *E*_*L*_ must be set to predetermined values, or additional experimental data are needed in the same recorded cell to constrain them, as discussed next.

To constrain *E*_*H*_, we simulated a tail current experiment that starts from a holding potential at which the H-current is activated (*V*_*T*,hold_ = −115 [mV]) and then steps to four potentials bracketing the presumed value of *E*_*H*_ (Fig. 1F). Note that the resulting current traces switch from convex to concave as the reversal potential value is crossed from below. This eliminates one of the remaining near-zero singular values (Fig. 1C, blue trace; see sec. S.3 for the corresponding analytical result). However, this adds nine parameters to the estimation problem: the steady-state activation at the holding potential (*h*_∞_(*V*_*T*,hold_)), and at each voltage step (*h*_∞_(*V*_*T,i*_), *i* = 1, …, 4), as well as the corresponding time constants (*τ*_*h*_(*V*_*T,i*_)).

To eliminate the last source of degeneracy, we simulated current injection after pharmacological block of the H-current, leaving a passive model (Fig. 1G). This current clamp experiment constrains *E*_*L*_ and removes the last near-zero singular value (Fig. 1C, red trace; see sec. S.4 for the corresponding analytical result). The final estimated parameters yield a unique solution close to the true parameter values (Table S2). The addition of noise to this model does not change qualitatively the results (Fig. S3).

### 2.2. H-current parameter identifiability in cell-attached configuration

Next, we analyzed the H-current of CA1 pyramidal cells characterized through cell-attached patch-clamp recordings [19, Fig. 1]. In that study, the cell-attached patch contained only H channels. Consequently, the model presented in eq. 15 does not include a leak current (see also [23] for a similar model in cerebellar Purkinje cells). Moreover, the selected ionic composition of the recording solution in the pipette resulted in the H-current reversal potential *E*_*H*_ being approximately equal to 0 [mV] [19]. Somatic and dendritic membrane patches had different numbers of H channels, resulting in different current magnitudes and noise levels (compare the y-axis in Fig. 2A and B; true parameters in Table S3). Additionally, the cell-attached recording configuration in [19] allowed to select a holding potential *V*_*C*,hold_ = −45 [mV] for which the H conductance was closed (*h*_∞_(*V*_*C*,hold_) = 0), in contrast to the holding potential near rest for the LGMD intracellular recordings (*V*_*C*,hold_ = − 60 [mV]).

As for the LGMD model, we initially estimate the unknown parameters by fitting the model to the voltage clamp portion of the recording (i.e., from 150 to 1140 [ms] in Fig. 2A and B). There are seventeen parameters: the maximal conductance 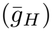, the reversal potential (*E*_*H*_), the steady-state activation at the holding potential (*h*_∞_(*V*_*C*,hold_)) and at each voltage step (*h*_∞_(*V*_*C,i*_), *i* = 1, …, 7), as well as the corresponding activation time constants (*τ*_*h*_(*V*_*C,i*_)). Like in the LGMD model, we fix the steady-state activation at the largest hyperpolarizing voltage step: *h*_∞_(*V*_*C*,7_) = 1 (*V*_*C*,7_ = − 125 [mV]).

The initial fit of the noise-free data (black dashed lines in Fig. 2A, B) yields one singular value close to zero (Fig. 2C, green). Application of the subset selection algorithm flags *E*_*H*_ as the least identifiable parameter, as in the LGMD fits. Dendritic recordings yielded similar results.

Surprisingly, and in contrast to LGMD model (Fig. S3), when measurement noise was included, the gap in the singular value spectrum was considerably reduced, resulting in a full-rank Jacobian (Fig. 2C, red). Thus, in this case, noise can mask the existence of an unconstrained model parameter. Analytical results show that this can be traced back to the fact that the CA1 H-current model lacks a leak current and that *h*_∞_(*V*_*C*,hold_) = 0 (sec. S.5). Nonetheless, the Jacobian remained poorly conditioned because the singular values span a large range.

To further improve parameter identifiability, the tail portion of the voltage clamp experiment was fitted, providing additional constraints on *E*_*H*_, as in the LGMD case. This added one parameter to the fitting procedure: the activation time constant at *V*_*C*,hold_. Yet, including this additional data decreased the range of Jacobian singular values, indicating improved parameter identifiability (Fig. 2C, blue; see sec. S.6 for the corresponding analytical result). The estimated steady-state activation at each voltage step (*h*_∞_(*V*_*C,i*_)) for somatic and dendritic recordings and their 95% confidence intervals, represented by error bars, demonstrate accurate parameter estimates despite the recording noise (Fig. 2D; sec. 4.2.5).

### 2.3. Impact of inactivation on parameter identifiability

A-type currents include both activation and inactivation terms. This example was taken from whole-cell recordings of Kenyon cells in honeybees (Fig. 3A; true parameters in Table S4) [20, Fig. 6].

**Fig 3.**
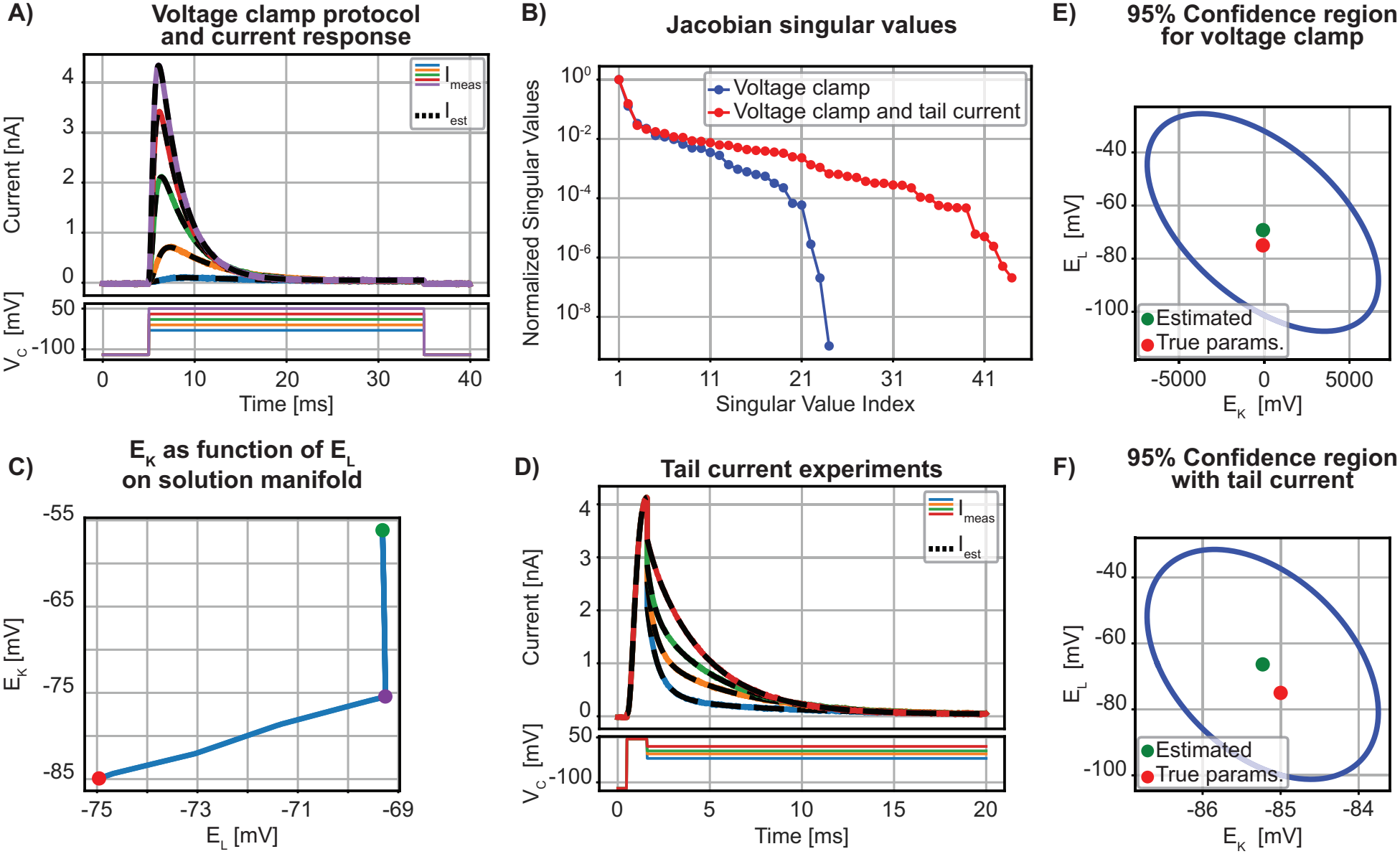
Parameter estimation and degeneracy for the A-current model. **(A)** Voltage clamp data used for parameter estimation and the corresponding model fits (colored and black dashed lines, respectively). **(B)** Normalized Jacobian singular values before and after adding tail current experiments. Tail current experiments improve the Jacobian condition number by constraining *E*_*K*_. **(C)** *E*_*K*_ as a function of *E*_*L*_ on the solution manifold obtained from voltage clamp experiments. Parameter degeneracy is due to an ill-conditioned estimation problem, despite a full-rank Jacobian. The three colored dots represent selected examples of parameter sets on the manifold that generate nearly identical voltage clamp outputs (Fig. S4). **(D)** Tail current traces used to constrain *E*_*K*_ and the corresponding model fits (colored and black dashed lines, respectively). **(E)** 95% confidence regions for *E*_*K*_ and *E*_*L*_ before and **(F)** after the inclusion of tail current experiments. **(E)** shows large uncertainty in *E*_*K*_, significantly reduced in **(F)**.

According to eq. 18, model-based multiplicative parameter degeneracy involves 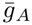, as well as the activation (*n*) and inactivation (*h*) terms. To eliminate this degeneracy, we assumed full activation (*n*_∞_(*V*_*C*,5_) = 1) at the largest voltage step (*V*_*C*,5_ = 50 [mV]) and full de-inactivation at the holding potential (*h*_∞_(*V*_*C*,hold_) = 1). The model contains twenty-four remaining parameters: the maximal conductances 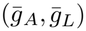, the reversal potentials (*E*_*K*_, *E*_*L*_), the steady-state activation at the holding potential (*n*_∞_(*V*_*C*,hold_)), the steady-state activation (*n*_∞_(*V*_*C,i*_), *i* = 1, …, 5) and inactivation (*h*_∞_(*V*_*C,i*_), *i* = 1, …, 5) at each voltage step (except as fixed above), as well as the corresponding time constants of activation (*τ*_*n*_(*V*_*C,i*_), *i* = 1, …, 5) and inactivation (*τ*_*h*_(*V*_*C,i*_), *i* = 1, …, 5). Estimation through nonlinear least-squares generated a successful fit to the measured data (Fig. 3A).

In contrast to the H-current LGMD model, the inclusion of an inactivation term results in a full-rank Jacobian (Fig. 3B; see sec. S.7 for the corresponding analytical result). However, the singular values are widely spread and therefore the Jacobian matrix has a high condition number [22], suggesting the existence of a manifold of solutions with negligible differences in least-squares error. Numerical continuation demonstrates a wide range of possible values for *E*_*K*_, showing that this parameter is poorly constrained (Fig. 3C, Fig. S4). This suggests that tail current experiments may improve parameter identifiability (Fig. 3D).

To simulate tail current experiments [24], we first held the membrane potential at a large hyperpolarizing voltage, followed by a brief depolarizing pulse to activate the A-current. Then, we stepped to four voltages (*V*_*T,i*_) selected (by trial and error) to best constrain *E*_*K*_. Note that the fast activation of the A-current and its inactivation during hyperpolarizing voltage steps limit the range of tail voltages that can provide information constraining *E*_*K*_. These additional experiments add twenty parameters to the estimation problem: the pulse steady-state activation and inactivation terms (*n*_∞_(*V*_*T*,pulse_), *h*_∞_(*V*_*T*,pulse_)), their corresponding time constants (*τ*_*n*_(*V*_*T*,pulse_), *τ*_*h*_(*V*_*T*,pulse_)), terms at each voltage step for the steady-state activation (*n*_∞_(*V*_*T,i*_), *i* = 1, …, 4) and inactivation (*h*_∞_(*V*_*T,i*_), *i* = 1, …, 4), as well as the corresponding time constants of activation (*τ*_*n*_(*V*_*T,i*_), *i* = 1, …, 4) and inactivation (*τ*_*h*_(*V*_*T,i*_), *i* = 1, …, 4).

The effect of additional tail current data can also be observed in the 95% confidence regions (see also sec. 2.5). The voltage clamp data alone poorly constrained *E*_*K*_ (note the horizontal axis range in Fig. 3E). Incorporating the tail current experiment significantly narrows the confidence region in the *E*_*K*_ direction (Fig. 3F).

### 2.4. Parameter identifiability of the Hodgkin-Huxley model

Next, we studied the action potential generating currents in the Hodgkin-Huxley squid giant axon model (true parameters in Table S5; gating kinetics in sec. S.8) [1]. The sodium current model eq. 22 has thirty-four parameters: the maximal conductances 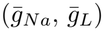, reversal potentials (*E*_*Na*_, *E*_*L*_), the steady-state activation and inactivation at the holding potential (*m*_∞_(*V*_*C*,hold_), *h*_∞_(*V*_*C*,hold_)) and at each voltage step (*m*_∞_(*V*_*C,i*_), *h*_∞_(*V*_*C,i*_), *i* = 1, …, 7), as well as the corresponding activation and inactivation time constants (*τ*_*m*_(*V*_*C,i*_), *τ*_*h*_(*V*_*C,i*_), *i* = 1, …, 7). To eliminate model-based multiplicative parameter degeneracy, we fix *m*_∞_(*V*_*C*,7_) = 1 at the largest depolarizing voltage step (*V*_*C*,7_ = 62 [mV]). And as in [1, Table 2], we set *h*_∞_(*V*_*C*,hold_) = 0.6 at the holding potential (*V*_*C*,hold_ = − 65 [mV]). This leaves thirty-two parameters to fit and results in a full rank Jacobian (sec. S.9).

The potassium current model eq. 25 has nineteen parameters: the maximal conductances 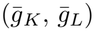, reversal potentials (*E*_*K*_, *E*_*L*_), the steady-state activation at the holding potential (*n*_∞_(*V*_*C*,hold_)) and at each voltage step (*n*_∞_(*V*_*C,i*_), *i* = 1, …, 7), as well as the corresponding activation time constants (*τ*_*n*_(*V*_*C,i*_)). To eliminate multiplicative parameter degeneracy we set *n*_∞_(*V*_*C*,7_) = 1 (*V*_*C*,7_ = 62 [mV]) leaving eighteen parameters to fit.

Parameter estimation produced successful fits to the measured sodium and potassium currents (black dashed lines in Fig. 4A, B). Despite the similarity between the potassium and H-current models, both being described solely by activation dynamics, their Jacobian singular value spectra differ. Based on the H-current model results, we expected two near-zero normalized singular values, with *E*_*K*_ and *E*_*L*_ being the least identifiable, as in Fig. 1C. Surprisingly, no singular value approaches zero, and the residual Jacobian is full rank (Fig. 4E, blue; see sec. S.9 for the corresponding analytical result). This difference originates from the fourth-order activation term in the potassium current model, which improves parameter identifiability. Indeed, replacing the fourth-order activation by a first-order one leads to Jacobian rank deficiency, like in the H-current model and in agreement with mathematical results (Fig. 4E, red trace, and sec. S.1). The same holds for the sodium current, where third-order activation yields a narrower singular value spectrum than a model with first-order activation (Fig. 4D).

**Fig 4.**
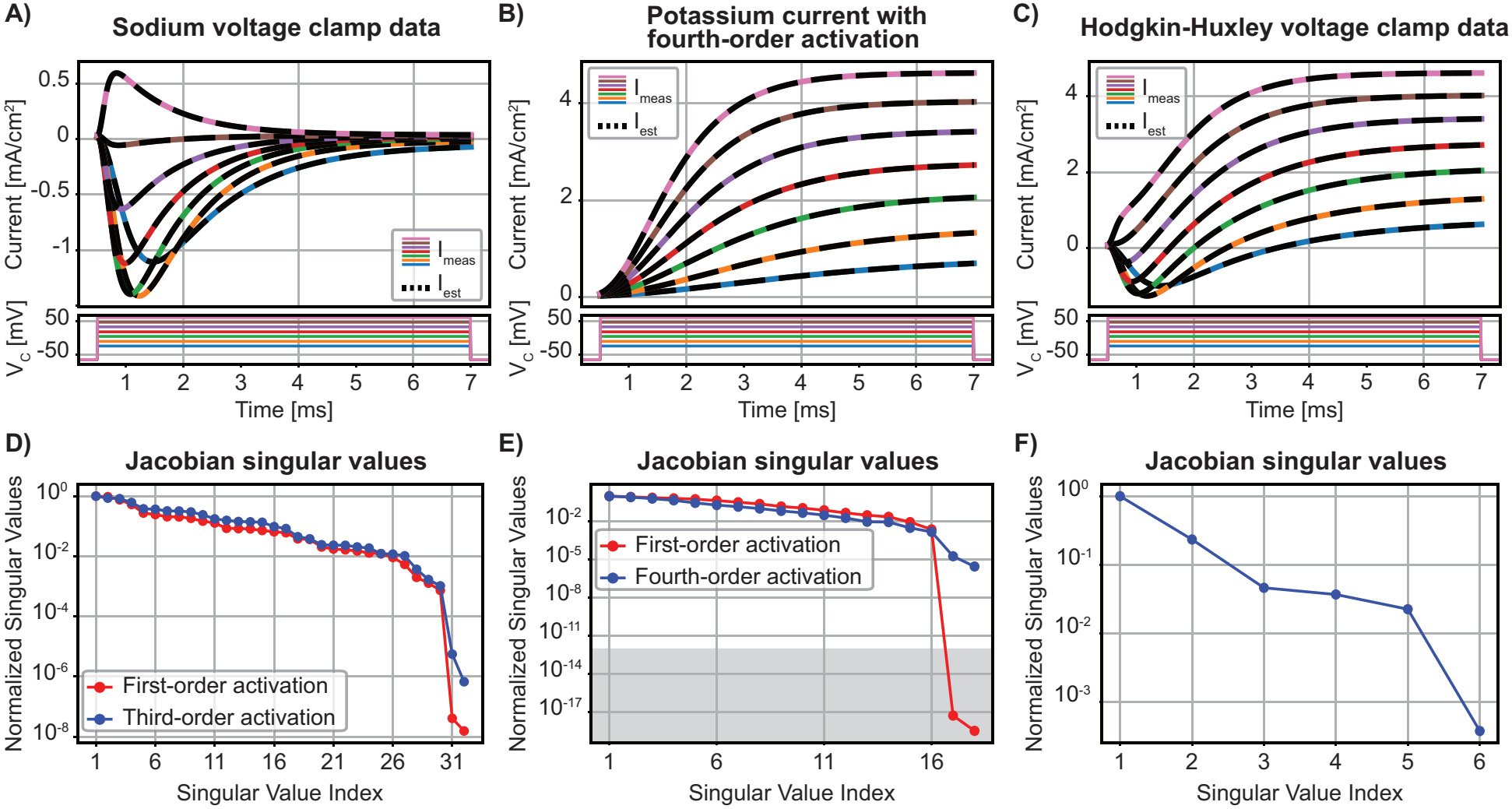
Parameter estimation and degeneracy in the Hodgkin-Huxley model. **(A)** Sodium current voltage clamp data and the corresponding model fits (colored and black dashed lines, respectively). **(B)** Potassium current voltage clamp data and the corresponding model fits presented as in A. **(C)** HH voltage clamp data including both currents and the corresponding model fits presented as in A. **(D)** Normalized singular values for the sodium model with first-order (red) vs. third-order activation (blue). **(E)** Normalized singular values for the potassium model with first-order (red) vs. fourth-order activation (blue). Singular values that fall within the gray region, below the threshold of 10^−12^, are considered to be equal to zero. **(F)** Normalized singular values for the full HH model used to estimate the maximal conductances and reversal potentials.

Next, we used the full HH model eq. 28 to estimate peak conductances and reversal potentials from voltage clamp data obtained without separating pharmacologically the potassium and sodium currents (Fig. 4C; adapted from [25, Fig. 3b]). The six parameters are: 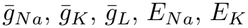, and *E*_*L*_. Inspection of the singular values indicates a full rank Jacobian and thus a unique solution (Fig. 4F). Moreover, the small spread of the singular values indicates that the parameters are well constrained. As a result, no solution manifold is detected.

### 2.5. Parameter correlations and confidence regions

An advantage of estimating all parameters simultaneously is the ability to analyze their relationships. To identify groups of highly correlated parameters, we performed hierarchical clustering on pairwise parameter correlations, derived from the estimated Hessian at the solution (Fig. 5A; sec. 4.2.5). From these pairwise correlation coefficients, we derived 95% confidence regions, providing further insight into the structure of parameter uncertainty. For example, in the CA1 pyramidal cell H-current model, the highly correlated parameters *E*_*H*_ and 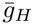 (orange box in Fig. 5A) produce a narrow, elongated confidence ellipse (Fig. 5B). Although this is expected for highly correlated parameters, its tilted orientation reveals a hidden direction along which the uncertainty is greatest. This information cannot be obtained from individual confidence intervals or from correlation coefficients alone.

**Fig 5.**
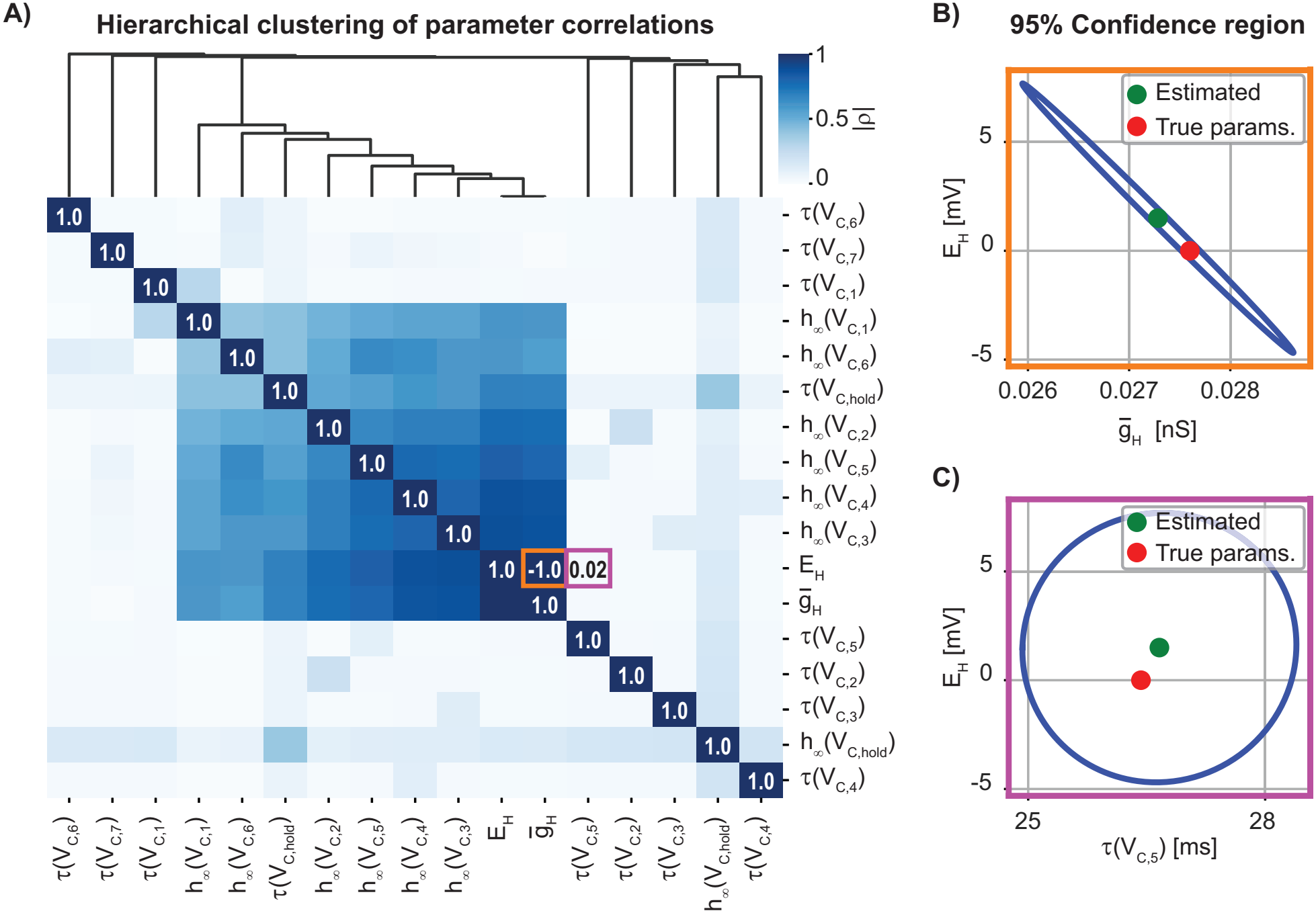
Parameter correlations and confidence regions for the CA1 pyramidal cell H-current model. **(A)** Hierarchical clustering of the correlation matrix reveals relationships among parameters and identifies groups of highly correlated parameters. **(B, C)** Examples of 95% confidence regions for selected parameter pairs, marked in orange and purple in (A). (B) corresponds to the orange box in (A), where the selected parameters are strongly correlated. The narrow and elongated ellipse reveals a direction of uncertainty in parameter space that is not apparent from the individual confidence intervals alone. (C) corresponds to the purple box in (A), where the selected parameters are weakly correlated, resulting in a nearly circular confidence region and nearly independent parameter uncertainties.

In contrast, uncorrelated parameters (e.g., purple box in Fig. 5A) produce a nearly circular confidence region (Fig. 5C). In this case, the uncertainties of the two selected parameters are largely independent, and there is no preferred direction along which changes in one parameter can be compensated by changes in the other.

Moderately strong correlations are also present among a subset of parameters consisting mainly of steady-state activation values (Fig. 5A). The correlation structure is different for the H-current model associated with LGMD intracellular voltage clamp data, with the strongest correlations among pairs of variables rather than in a common block (Fig. S5). In the same model, the assumption that *h*_∞_(*V*_*C*,5_) = 1 leads to a systematic error when estimating 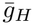 (Fig. S6A; see also sec. 2.1 and sec. S.2). Other parameters remain unaffected (Fig. S6B). The correlation structure for the A-type current presents features of both H-current models, exhibiting a large block of correlated parameters and several smaller ones (Fig. S7). Similar features were also seen in the correlation structure of the HH sodium and potassium currents (Figs. S8, S9).

We also observed that across all models, the maximal conductance and reversal potential of an ionic current were consistently highly correlated, including the leak, when present (see red and orange boxes in Figs. 5A, S5, S7, S8, and S9).

## 3. Discussion

In this work, we identify parameter degeneracy in least-squares HH-type current models by computing the spectrum of singular values associated with the matrix consisting of the derivatives of the residuals with respect to the model parameters (i.e., the Jacobian matrix). We find that singular values typically span a broad range – a defining characteristic of numerically rank-deficient or ill-conditioned least-squares optimization problems [22]. In all cases, rank deficiencies in the Jacobian matrix observed numerically using the singular values are proven analytically [21] (secs. S.1, S.3 – S.7, and S.9). Consequently, our results are generally applicable – irrespective of the particular current studied – under the stated experimental conditions and for HH-type current models whose functional form conforms to the theorem statements provided in the Supporting Information.

### A toolkit to identify parameter degeneracy and solution manifolds

The comprehensive approach developed in sec. 2 is summarized in Fig. S10. In this context, while the singular value spectrum associated with a least-squares problem is a well-known tool [22, 26, 27], to our knowledge it has not been applied to the fitting of HH-type current models.

For numerically rank-deficient problems such as the H-current LGMD model, subset selection identifies the least identifiable parameters from the Jacobian SVD. This allows either to fix these parameters, or to design further experiments constraining them. The use of subset selection is new in our settings, but has been applied in other contexts when insufficient data is available to constrain model parameters [28]. An alternative is to assign Bayesian priors [28] (in the context of HH-type models, see [29]). However, this is equivalent to fixing the parameters to the most likely prior in numerically rank-deficient problems, since no additional information is available to constrain them (besides the prior).

Importantly, our results show that one can also use the SVD of the Jacobian to discover the directions along which to trace the manifold of equivalent parameter solutions for the least squares minimization problem. Continuation methods have been studied in different contexts, see, e.g., [30], [31], [32], but their use to reveal the solution manifold of an unconstrained nonlinear least squares problem such as eq. 30 is new.

### Biophysical implications

For the LGMD H- and the A-current models, parameter manifold tracing shows that several parameters need to change in unison to maintain the same solution to the least squares problem (though not all change; Figs. 1E, 3C, S1, S2). In contrast, fits of the HH-current models and of the CA1 H-current model after adding tail current experiments did not reveal parameter degeneracy. The experimental conditions in these two cases were close to ideal, with the isolation of a single channel type, and a large space clamp area allowing for nearly noise-free recordings, respectively. Nonetheless, even under nearly ideal conditions, our results show that not all parameters are equally well constrained. For example, in the CA1 H-current model, the activation curves are tightly constrained, confirming a reliable and yet subtle difference between soma and dendrites (Fig. 2D). In contrast, the reversal potential could vary by as much as *±* 5 [mV], provided the maximal conductance changes accordingly (Fig. 5B; for the HH model, see Figs. S8 and S9).

Interestingly, the same correlation between peak conductance and reversal potential was found in all models, irrespective of current type. This can be understood by recalling that the peak conductance multiplies the driving force, i.e., the difference between the clamp potential and the reversal potential. A peak conductance increase can thus be compensated by a concurrent decrease in driving force, or equivalently a concurrent change in the reversal potential since the driving force keeps the same sign in most voltage clamp experiments. To our knowledge, this correlation has not been reported earlier, but reversal potentials have rarely been considered as parameters in voltage clamp fits. Their cell-to-cell variability has been reported when measured [1, 33], as expected from the complex mechanisms maintaining their values through ion pumps and exchangers [34–36]. Asserting whether peak conductance and reversal potentials change in parallel from one experiment to the next – as predicted here – would require simultaneous measurements *in vivo*.

### Relation to previous work

Our results complement and extend earlier work, some of which has already been described in the introduction. Additionally, model-based multiplicative parameter degeneracy was analyzed in [37, 38]. Simultaneous fitting of multiple parameters has also been treated in [39]. The implications of parameter uncertainty on neuronal excitability in the squid giant axon HH model have been studied in [33, 40]. Note that our parameter manifold tracing technique differs substantially in its complexity from the one considered in [41]. There, the dependency of a few measured characteristics of the model output as a function of the model parameters was considered and fitted to experimental data. This approach allowed the authors to apply the implicit function theorem to determine a local manifold of parameters on which the measured characteristics remain constant. Since the number *k* = 2 of measured characteristics in [41] is substantially smaller than the total number *n* of parameters, the implicit function theorem guarantees that *n* − *k* parameters can be written as a function of *k* suitably chosen ones (under standard assumptions, e.g., differentiability of the measured characteristics with respect to the model parameters). In contrast, we apply parameter continuation directly to the *m >> n* measurements rather than to *k < n* measured characteristics derived from the measurements. In the terminology of [42], our results show that HH-type models like the H-current LGMD model are structurally unidentifiable when only voltage clamp data is available. However, our results go far beyond those of [42] by developing comprehensive methods that reveal when HH-type models are structurally unidentifiable and that fully characterize their parameter degeneracy.

### Future directions and summary

It should be straightforward to generalize our approach to conductance-based models with different gating schemes, e.g., when multiple gating variables are not independent of each other [43, Chap. 18]. Fits of the membrane potential in compartmental models to current clamp experiments in single neurons are also expected to be amenable to a similar analysis. Specifically, there exist well-documented methods to fit simultaneously the parameters of ordinary differential equations that could be used in conjunction with our methods [44]. In summary, the methods presented here open the possibility of better understanding how robust neuronal computations are to the biophysical parameters of single neurons, an important topic in neuroscience [3].

## 4. Models and methods

### 4.1 Hodgkin-Huxley-type compartmental neural models

The general form of an HH-type model is

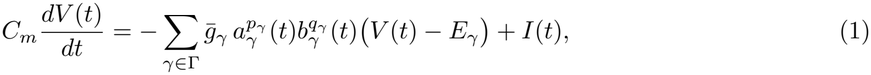

where *C*_*m*_ is the membrane capacitance, Γ is a set of channels considered, 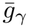 and *E*_*γ*_ are the maximal conductance and reversal potential of ionic current *γ*, respectively. The variable *I*(*t*) is the recorded or injected current. The variables *a*_*γ*_ and *b*_*γ*_ are voltage-dependent gating variables with corresponding exponents *p*_*γ*_ and *q*_*γ*_ satisfying first order linear ordinary differential equations:

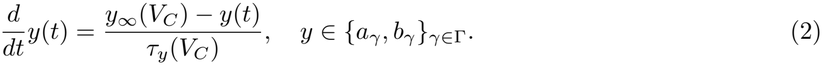

By modifying the conductance parameters and gating dynamics, eq. 1 and eq. 2 can describe a wide range of ionic currents.

During a voltage clamp experiment, the membrane potential is held at a prescribed holding potential (*V*_*C*,hold_) and then instantaneously stepped to a command voltage (*V*_*C*_) at time *t*_*V*,0_. Therefore, d*V/*d*t* = 0, and eq. 1 reduces to

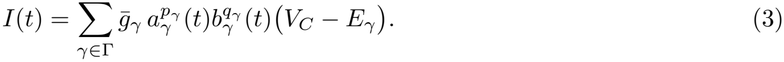

#### 4.1.1. Hyperpolarization-activated ionic current in the LGMD neuron

This model follows the form presented in eq. 1 – eq. 3 with Γ = {*L, H*}, *p*_*L*_ = *q*_*L*_ = 0, *p*_*H*_ = 1, *q*_*H*_ = 0, and *a*_*H*_ = *h*. The H-current steady-state activation and time constant functions are given by

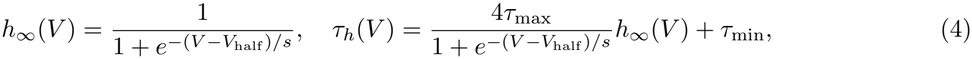

with true parameters specified in Table S1.

##### Voltage clamp experiment

Solving eq. 2 analytically for *y* = *h* shows that *h*(*t*) relaxes exponentially starting at time *t*_*V*,0_ from its value at the holding potential, *h*_∞_(*V*_*C*,hold_), toward its steady-state value at the command voltage, *h*_∞_(*V*_*C*_). Substitution in eq. 3 gives the analytical expression used to simulate the measured current:

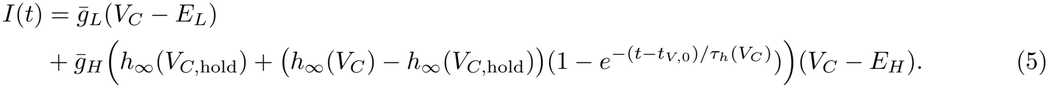

The holding membrane potential is set at *V*_*C*,hold_ = − 60 [mV], and stepped to one of the five command voltages

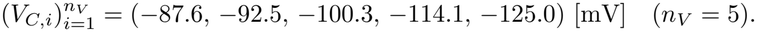

##### Parameters and objective function

For *n*_*V*_ voltage clamp experiments, we estimate

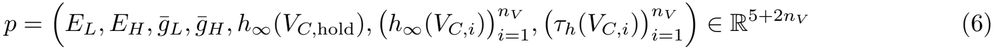

or when *h*_∞_(*V*_*C*,5_) = 1 (*V*_*C*,5_ = −125 [mV]),

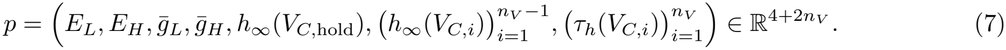

The parameter estimates are minimizers of the squared error between the measured current from the voltage clamp experiment and the current predicted by the model eq. 5 at times *t*_*V,j*_, *j* = 1, …, *m*_*V*_ . Therefore, the objective function is

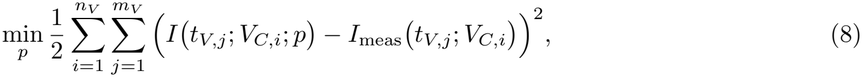

where the parameters *p* are given by eq. 6 or eq. 7, *I (t*_*V,j*_; *V*_*C,i*_; *p)* is the fu nction eq. 5 estimated at times *t*_*V,j*_ ∈ [0.5, 3.5) [s], with *V*_*C*_ = *V*_*C,i*_ and *t*_*V*,0_ = 0.5 [s], while *I*_meas_ *(t*_*V,j*_ ; *V*_*C,i*_) are the measured currents.

##### Tail current experiment

The membrane potential is held at a hyperpolarizing voltage, *V*_*T*,hold_ = −115 [mV], to activate the H-current. We then take four voltage steps that bracket the expected value of *E*_*H*_,

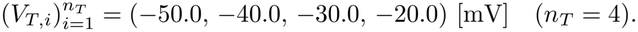

##### Parameters and objective function

Including tail current data introduces 1 + 2*n*_*T*_ parameters to the estimation problem. Consequently, eq. 6 and eq. 7 are replaced by

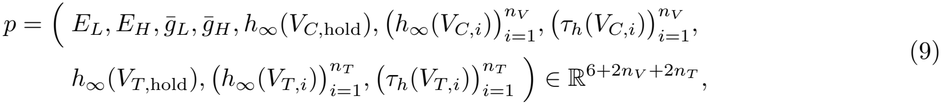

and

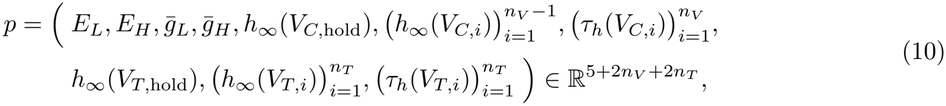

respectively.

The objective function is

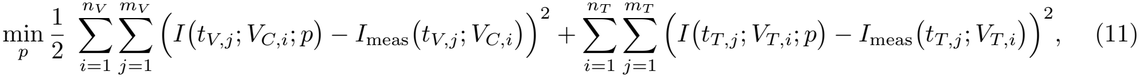

where the parameters *p* are given by eq. 9 or eq. 10. The terms in the first double sum are given as before, and in the second double sum *I t*_*T,j*_; *V*_*T,i*_; *p* is the function eq. 5 evaluated at times *t*_*T,j*_ ∈ [3.5, 4.5) [s], with *V*_*C*,hold_, *V*_*C*_ replaced by *V*_*T*,hold_, *V*_*T,i*_, respectively, and *t*_*T*,0_ = 3.5 [s]. *I*_meas_ *t*_*T,j*_; *V*_*T,i*_ are the measured tail currents.

##### Current clamp experiment in a passive model

A constant current injection, *I*_*C*_, is initiated at time *t*_*C*,0_ and continues until *t*_*C,f*_ . Following eq. 1, the membrane voltage dynamics is described by

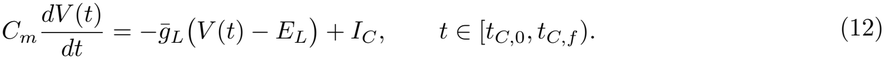

Using the method of integrating factors, the solution of eq. 12 is

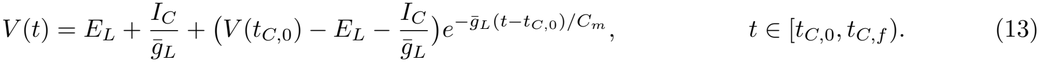

In the interval [0, *t*_*C*,0_) no current is injected (*I*_*C*_ = 0) and the voltage remains as the initial condition *V* (0) = *E*_*L*_. As a result, *V* (*t*_*C*,0_) = *E*_*L*_. We simulate three constant current injections

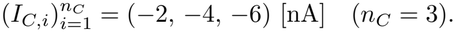

Throughout the simulation, the membrane capacitance is fixed at *C*_*m*_ = 0.0018 [*µ*F].

##### Parameters and objective function

Unlike the tail current experiment, the current clamp experiment does not introduce additional unknown parameters. As a result, when *n*_*V*_ voltage clamp experiments, *n*_*T*_ tail current experiments, and *n*_*C*_ current clamp experiments are used simultaneously, we still estimate the parameters in eq. 9 or eq. 10 but now as the solution of

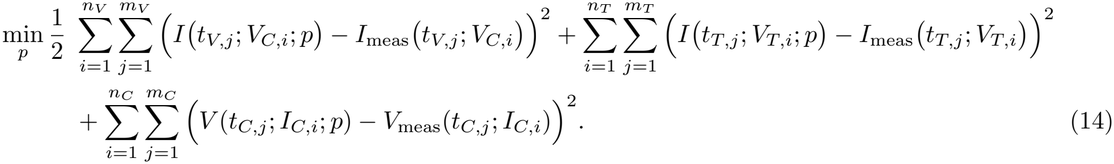

In the third double sum, *V* (*t*_*C,j*_; *I*_*C,i*_; *p*) is computed from eq. 13 with *t*_*C*,0_ = 0.5 [s], *t*_*C,j*_ ∈ [0.5, 3.5) [s], and *I*_*C*_ = *I*_*C,i*_ (*i* = 1, 2, 3). *V*_meas_(*t*_*C,j*_; *I*_*C,i*_) are the measured voltages.

#### 4.1.2 Hyperpolarization-activated current in CA1 pyramidal cells

The model is similar to that of the H-current in the LGMD, but uses the true parameters specified in Table S3.

##### Voltage clamp experiment

After removing the leak current in eq. 5, the synthetic data are generated using

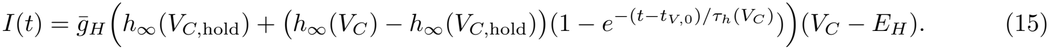

The voltage clamp protocol is adopted from [19, Fig. 1]. The holding potential is *V*_*C*,hold_ = 45 [mV], followed by seven voltage steps for somatic and dendritic recordings, respectively,

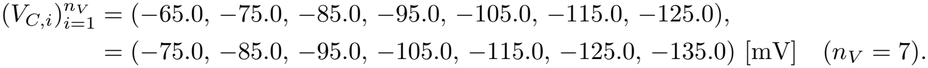

##### Parameters and objective function

For *n*_*V*_ voltage clamp experiments, we estimate the parameters

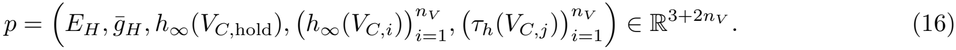

The parameter estimates are obtained by solving a modified version of eq. 8 with model eq. 5 replaced by eq. 15. As for the LGMD model, we fix 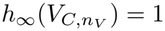 and remove it from the list of parameters *p* in eq. 16, to avoid model-based degeneracy due to multiplicative combination of 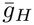 and *h*_∞_ in eq. 15.

##### Tail current experiment

Following [19, Fig. 1], we simulate in the same voltage clamp protocol the step back from each command voltage (*V*_*C,i*_) at *t*_*T*,0_ = 1140 [ms] to the holding potential (*V*_*C*,hold_).

##### Parameters and objective function

This introduces one additional parameter, the activation time constant at the holding potential *τ*_*h*_(*V*_*C*,hold_). We thus estimate 3 + 2*n*_*V*_ parameters using a modified version of eq. 11

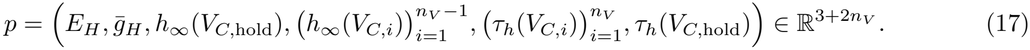

#### 4.1.3 A-Type potassium current

The model follows the form presented in eq. 1 – eq. 3 with Γ = {*L, A*}, *p*_*L*_ = *q*_*L*_ = 0, *p*_*A*_ = 3, *q*_*A*_ = 1, *a*_*A*_ = *n*, and *b*_*A*_ = *h*. The voltage-dependent steady-state values of the gating variables and time constants are

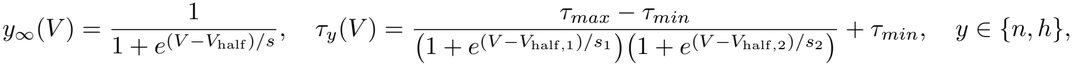

with true parameters specified in Table S4 [20].

##### Voltage clamp experiment

Similar to the analytical solution provided in eq. 5, the current corresponding to each command voltage *V*_*C*_ can be written as follows

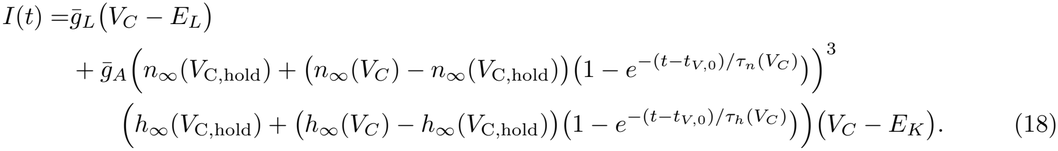

Note that we use *E*_*K*_ to denote the A-type current reversal potential, as this ion channel is predominantly permeable to potassium ions (*K*^+^). The voltage clamp protocol is adapted from [20, Fig. 6]. The membrane potential is held at a large hyperpolarized voltage *V*_*C*,hold_ = − 120 [mV], and then stepped to five command voltages

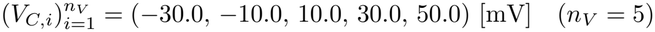

for 30 [ms]. We then step back to the holding potential to complete the voltage clamp simulation.

##### Parameters and objective function

For *n*_*V*_ voltage clamp experiments, we estimate the unknown parameters

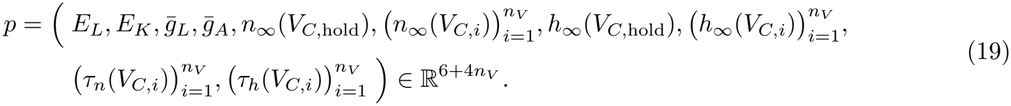

Fixing *h*_∞_(*V*_*C*,hold_) = 1 and *n*_∞_(*V*_*C*,5_) = 1 to eliminate multiplicative model-based degeneracy, results in

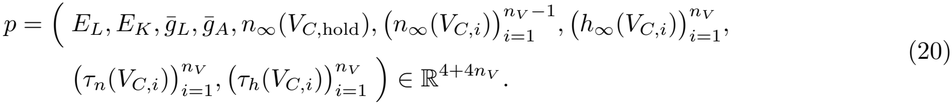

We use a modified version of eq. 8 to estimate the parameters. In these simulations *t*_*V*,0_ = 5.0 [ms], and the model is evaluated at times *t*_*V,j*_ ∈ [5.0, 35.0) [ms].

##### Tail current experiment

We adapt a protocol from [24, Fig. 4]. The membrane potential is held at a hyperpolarized voltage (*V*_*T*,hold_ = −120 [mV]) to ensure the A-current is fully de-inactivated. Next, the A-current is activated by briefly stepping to a depolarizing voltage *V*_*T*,pulse_ = 45 [mV] for 1.1 [ms]. We then immediately step to the four selected voltages

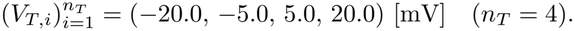

##### Parameters and objective function

Including the tail current experiment in the estimation problem introduces 4 + 4*n*_*T*_ additional parameters,

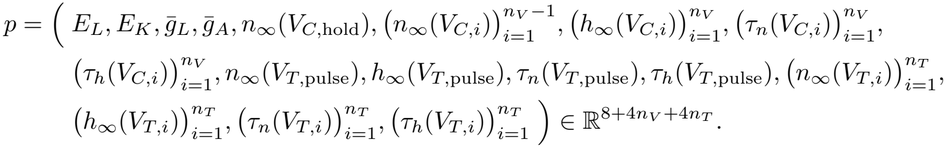

The parameters are estimated by solving the least squares problem

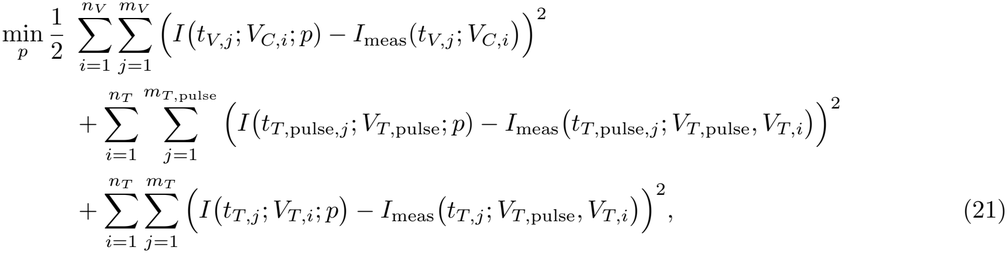

where the terms in the first double sum are given as before. In the second double sum, *I (t*_*T*,pulse,*j*_; *V*_*T*,pulse_; *p)* is the function eq. 18 evaluated at times *t*_*T*,pulse,*j*_ ∈ [0.5, 1.6) [ms] with *V* _*C*,hold_, *V*_*C*_ replaced by *V*_*T*,hold_, *V*_*T*,pulse_, respectively, and *t*_*T*,pulse,0_ = 0.5 [ms]. *I*_meas_ *(t*_*T*,pulse,*j*_ ; *V*_*T*,pulse_, *V*_*T,i*_) are the measured activation pulse currents of the experiment where the A-current is activated by briefly stepping to a depolarizing voltage *V*_*T*,pulse_ for 1.1 [ms] and then immediately stepped to the selected voltage *V*_*T,i*_. In the third double sum, *I (t*_*T,j*_; *V*_*T,i*_; *p)* is the function eq. 18 evaluated at t imes *t*_*T,j*_ ∈ [1.6, 20 .0) [ms] with *V*_*C*,hold_, *V*_*C*_ replaced by *V*_*T*,pulse_, *V*_*T,i*_, respectively, and *t*_*T*,0_ = 1.6 [ms]. *I*_meas_ *(t*_*T,j*_ ; *V*_*T*,pulse_, *V*_*T,i*_) are the measured tail currents after the voltage clamp pulse to *V*_*T*,pulse_.

Note that if the activating pulse voltage (*V*_*T*,pulse_) is selected from one of the command voltages (*V*_*C,i*_) introduced in the voltage clamp experiment, the four corresponding parameters, *n*_∞_(*V*_*T*,pulse_), *h*_∞_(*V*_*T*,pulse_), *τ*_*n*_(*V*_*T*,pulse_), and *τ*_*h*_(*V*_*T*,pulse_), are already included in the voltage clamp estimation problem. As a result, the problem can be simplified by removing the second sum in eq. 21 and estimating the 4 + 4*n*_*V*_ + 4*n*_*T*_ parameters.

#### 4.1.4. Hodgkin and Huxley model

##### Sodium current model

The model follows the form presented in eq. 1 – eq. 3 with Γ = {*L, Na*}, *p*_*L*_ = *q*_*L*_ = 0, *p*_*Na*_ = 3, *q*_*Na*_ = 1, *a*_*Na*_ = *m*, and *b*_*Na*_ = *h*. The true parameters are derived from the rate functions in eq. S1 and eq. S2.

##### Voltage clamp experiment

Using the analytical solutions for the gating variables as in eq. 5 gives

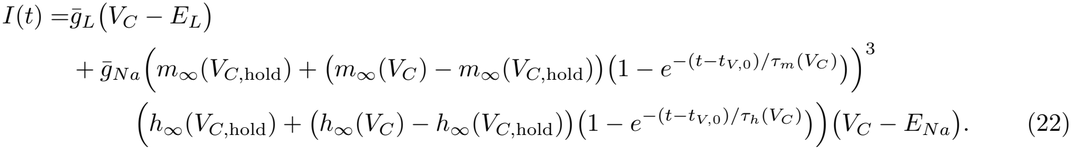

The voltage clamp protocol is adapted from the original Hodgkin and Huxley studies, where the squid giant axon was first held at its resting potential *V*_*C*,hold_ = −65 [mV], followed by seven voltage steps

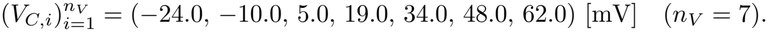

##### Parameters and objective function

Using the generated synthetic voltage clamp data, we estimate 6 + 4*n*_*V*_ parameters

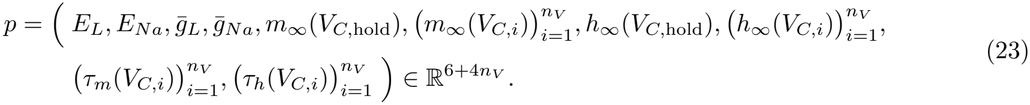

To resolve the model-based multiplicative parameter degeneracy in eq. 22, we set *h*_∞_(*V*_*C*,hold_) = 0.6 as in [1, Table 2], and *m*_∞_(*V*_*C*,7_) = 1 at the largest depolarizing command voltage (*V*_*C*,7_ = 62 [mV]). As a result, the remaining parameters are

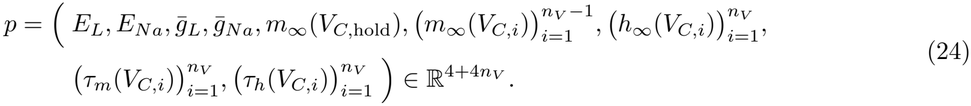

The parameter estimation problem is defined by minimizing eq. 8, where the parameters *p* are given by eq. 24, *I (t*_*V,j*_; *V*_*C,i*_; *p)* is the function eq. 22 evaluated at times *t*_*V,j*_ ∈ [0.5, 7.0) [ms], with *V*_*C*_ = *V*_*C,i*_ and *t*_*V*,0_ = 0.5 [ms].

##### Potassium current model

The model follows the form presented in eq. 1 – eq. 3 with Γ = {*L, K*}, *p*_*L*_ = *q*_*L*_ = 0, *p*_*K*_ = 4, *q*_*K*_ = 0, and *a*_*K*_ = *n*. The true parameters are derived from the rate functions in eq. S1 and eq. S3.

##### Voltage clamp experiment

Using the analytical solutions for the gating variables as in eq. 5 gives

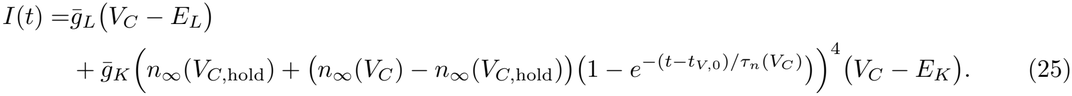

The voltage clamp protocol is identical to that of the sodium voltage clamp experiment.

##### Parameters and objective function

We estimate 5 + 2*n*_*V*_ parameters

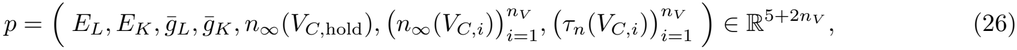

or after eliminating model-based multiplicative parameter degeneracy in eq. 25 by setting *n*_∞_(*V*_*C*,7_) = 1 (*V*_*C*,7_ = 62 [mV]),

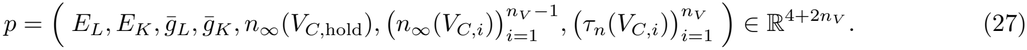

The parameter estimation problem is similar to eq. 8, where the parameters *p* are given by eq. 27, and *I (t*_*V,j*_; *V*_*C,i*_; *p)* is the function eq. 25.

##### Complete Hodgkin-Huxley model

The model combines the sodium and potassium currents since they were not separated pharmacologically, with true parameters given in Table S5.

##### Voltage clamp experiment

The resulting clamp current is given by

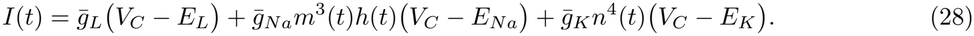

The voltage clamp protocol is identical to that of the sodium voltage clamp experiment.

##### Parameters and objective function

We estimate six parameters

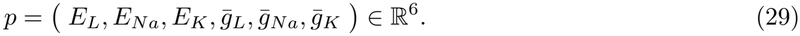

The estimation process is done by minimizing the least-squares error presented in eq. 8, where the parameters *p* are given by eq. 29, and *I (t*_*V,j*_; *V*_*C,i*_; *p)* is the function eq. 28.

#### 4.1.5. Noise determination

To simulate noisy recording data, we added Gaussian white noise to the model output. For each experimental data set, we measured the noise range from the steady-state portion of the voltage clamp recordings in the corresponding reference study. The standard deviation, *σ*_noise_, of the Gaussian white noise was then estimated by assuming that the noise range approximately corresponds to *±*3*σ*_noise_. The estimated *σ*_noise_ values are listed in Tables S1, S3 and S4.

### 4.2 Methods

The parameter estimation problems, such as eq. 8 or eq. 11, are nonlinear least squares problems of the type

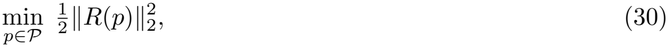

where *P* ⊂ ℝ ^*n*^ is a closed convex set of feasible parameters that encodes prior knowledge in the parameters, such as bounds parameters must obey, *R* : ℝ ^*n*^ → ℝ ^*m*^ is the residual function that describes the misfit between the output of the model with parameter *p* and the measurements, and 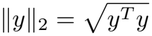 is the Euclidean norm. Often *P* = ℝ^*n*^ or *P* = {*p* ∈ ℝ^*n*^ : *a* ≤ *p* ≤ *b*}, where inequality constraints are understood componentwise.

More generally, we also consider weighted nonlinear least squares problems

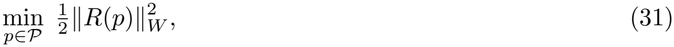

where ∥ *·* ∥_*W*_ is the weighted Euclidean norm

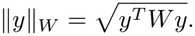

The symmetric positive definite weight matrix *W* is often chosen as the inverse of the covariance matrix of measurement errors [45, Chapter IV, Section 4-3], which in the linear case gives the estimate with the smallest covariance, see sec. 4.2.5.

Because *W* ∈ ℝ^*m×m*^ is symmetric positive definite, it can be factored as *W* = *LL*^*T*^ where *L* ∈ ℝ^*m×m*^ is nonsingular, using, e.g., the Cholesky decomposition [22, Sec. 4.2.3]. One can define the scaled residual function *L*^*T*^ *R* and write the weighted nonlinear least squares problem eq. 31 as an ordinary nonlinear least squares problem eq. 30 with *R* replaced by *L*^*T*^ *R*.

In addition to the explicit scaling of the residuals by a symmetric positive definite weight matrix *W*, see eq. 31, scaling of the parameters by a nonsingular matrix *D* ∈ ℝ;^*n×n*^ may be beneficial if parameters have vastly different scales. In that case, *R*(*p*) is replaced by *R*(*Dp*). Parameter scaling was not needed for the problems in this paper.

This section specifies the numerical methods for the solution of eq. 30, as well as numerical and statistical methods for the analysis of the computed solutions. An organogram summarizing how they are applied in sec. 2 is provided in Fig. S10.

#### 4.2.1 Nonlinear least squares solvers

The nonlinear least squares problems eq. 30 are solved iteratively using variations of the so-called Levenberg-Marquardt method [46, p. 227], [47, Section 6.2], [48, Section 10.3].

Our models and methods are implemented and executed in the Python programming environment. To make the reproducibility of our results easier, we use public domain nonlinear least squares solvers from SciPy (version 1.18.0) [49] and <monospace>lmfit</monospace> [50]. The <monospace>scipy.optimize.least_squares</monospace> function with the <monospace>method=‘lm’ </monospace> is a wrapper around the <monospace>MINPACK</monospace> [51] solver that implements the trust-region version of the Levenberg-Marquardt method in [52], and applies to unconstrained nonlinear least squares problems eq. 30 with *P* = ℝ^*n*^. The <monospace>scipy.optimize.least_squares</monospace> function with the <monospace>method=‘trf’ </monospace> is an implementation of the trust-region algorithm [53] and can handle bound-constrained nonlinear least squares problems eq. 30 with *P* = {*p* ∈ ℝ^*n*^ : *a* ≤ *p* ≤ *b*} . These two optimizers use analytical Jacobians, which we generate using automatic differentiation via the JAX Python library [54] with just-in-time (JIT) compilation.

In most cases, solving the unconstrained nonlinear least squares problem (i.e., *P* = ℝ^*n*^) using <monospace>scipy.optimize.least_squares</monospace> with <monospace>method=‘lm’</monospace> and with <monospace>method=‘trf’</monospace> gave similar results. However, in some cases, <monospace>method=‘lm’</monospace> converged to solutions that are not biophysically feasible. In those cases, we imposed bounds on the parameters, *P* = {*p* ∈ ℝ^*n*^ : *a* ≤ *p* ≤ *b*}, and used <monospace>method=‘trf’</monospace>. In a few of these cases, <monospace>method=‘trf’</monospace> did not converge to a solution, and we used <monospace>lmfit</monospace> with <monospace>method=‘leastsq’</monospace> to obtain an optimal solution that also satisfies the biophysical constraints.

#### 4.2.2 Numerical determination of Jacobian rank deficiency

The rank of a Jacobian *J*(*p*) can be obtained from its singular values *σ*_1_ ≥ … ≥ *σ*_*n*_ ≥ 0 as the largest integer *r* for which *σ*_*r*_ *>* 0 [22, p. 76]. Numerically, the rank *r* of the Jacobian can be estimated as the largest integer *r* for which the normalized singular value is above a threshold, *σ*_*r*_*/σ*_1_ ≥ *ϵ*, where *ϵ* is related to the machine precision used to compute the singular values [22, p. 275/276]. We selected *ϵ* = 10^−12^ as an appropriate cutoff based on the use of double-precision arithmetic and in combination with the rank results derived in [21].

#### 4.2.3. Subset Selection algorithm

As reported in Section 2, several voltage clamp experiments lead to nonlinear least squares problems eq. 30 with rank-deficient Jacobians, even though the number of measurements *m* is (much) larger than the number of parameters *n*. More specifically, the Jacobian *J*(*p*) ∈ ℝ^*m×n*^ of the residual function *R* has constant rank *r < n* for a wide range of parameters *p*. These rank results have been proven in [21] and have been summarized in sections S.1, S.3 – S.7, and S.9.

If the Jacobian of a nonlinear least squares problem eq. 30 has constant rank *r < n* for all parameters in a neighborhood of a solution *p*_*_ of eq. 30, then the constant rank theorem (see, e.g., [55, Theorem 4.12] or [56, Theorem 2.4.6]) implies the existence of a local manifold in parameter space containing *p*_*_ so that all points in this manifold are solutions of eq. 30.

So-called subset selection uses the Jacobian *J*(*p*) to indicate which *n* − *r* parameters can be fixed to remove this non-uniqueness. We call these parameters ‘least identifiable’. Our subset selection is from [22, Sec. 5.5.7] and is fully specified in Algorithm 1 below. Numerically, the algorithm selects *r* columns of the Jacobian *J*(*p*) that are sufficiently linearly independent, and so that the solution of the linear least squares problem with matrix given by these *r* columns results in a linear least squares residual that is close to the linear least squares residual computed with the full matrix.

##### Algorithm 1

Subset Selection

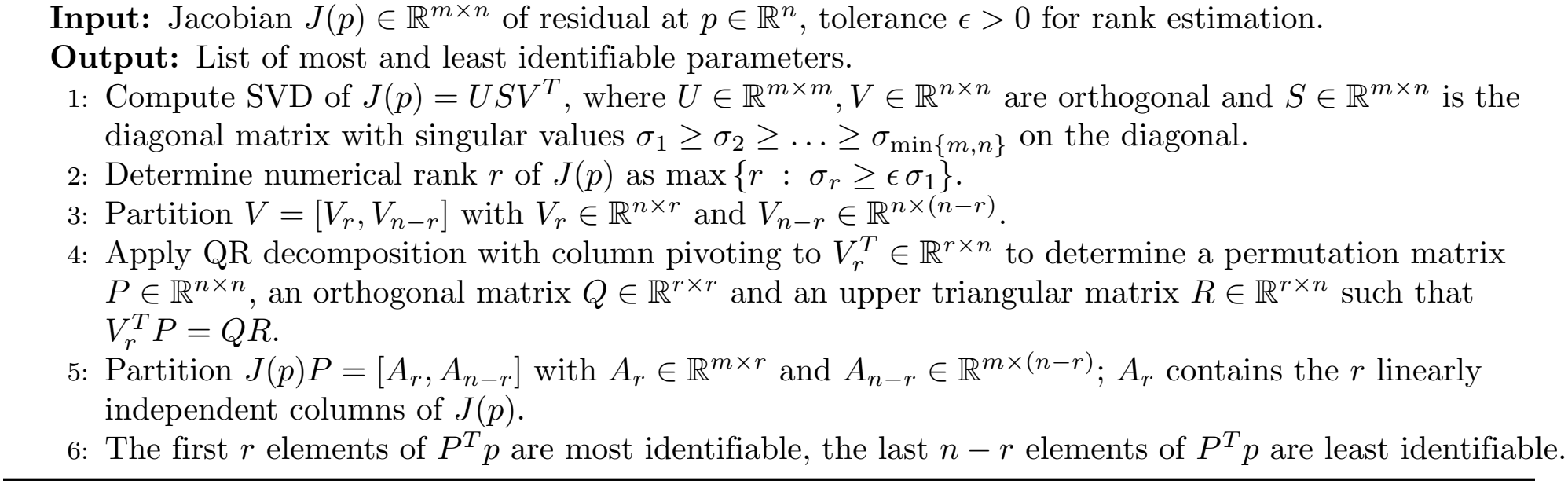

#### 4.2.4. Parameter continuation

As mentioned in the previous section 4.2.3, if the Jacobian *J*(*p*) ∈ ℝ ^*m×n*^ of an unconstrained (i.e., *P* = ℝ ^*n*^) nonlinear least squares problem eq. 30 has constant rank *r < n* for all parameters *p* in a neighborhood of a solution *p*_*_ of eq. 30, then the constant rank theorem implies the existence of a local smooth manifold in parameter space containing *p*_*_ so that all points in this manifold are solutions of eq. 30. More precisely, there exists a *q*_*_ ∈ ℝ ^*n*−*r*^ and a smooth function *p* : ℝ ^*n*−*r*^ → ℝ ^*n*^ defined in a neighborhood of *q*_*_ such that *p*(*q*_*_) = *p*_*_, *R*(*p*(*q*)) = *R*(*p*_*_), and the Jacobian *p*^*′*^(*q*) ∈ ℝ ^*n×*(*n*−*r*)^ of this function has rank *n* − *r* for all *q* ∈ ℝ ^*n*−*r*^ in a neighborhood of *q*_*_. Because *R* : ℝ ^*n*^ → ℝ ^*m*^ with *m > n*, the fact that *R*(*p*(*q*)) = *R*(*p*_*_) is constant for all *q* ∈ ℝ ^*n*−*r*^ in a neighborhood of *q*_*_ is a remarkable property that holds because the Jacobian of *R* has constant rank *r < n* in a neighborhood of *p*_*_.

In our setting, we know the local existence of a smooth function *p* : ℝ ^*n*−*r*^ → ℝ ^*n*^ such that *p*(*q*_*_) = *p*_*_, *R*(*p*(*q*)) = *R*(*p*_*_), and the Jacobian *p*^*′*^(*q*) ∈ ℝ ^*n×*(*n*−*r*)^ of this function at *q* has rank *n* − *r* for all *q* ∈ ℝ ^*n*−*r*^ in a neighborhood of *q*_*_, but this function is not known analytically. However, it can be approximated numerically using a continuation method. We describe our approach next.

Given *p*(*q*_*_) = *p*_*_, the Jacobian *p*^*′*^(*q*_*_) ∈ ℝ ^*n×*(*n*−*r*)^ of this function at *q*_*_ must satisfy *J*(*p*_*_)*p*^*′*^(*q*_*_) = 0 because *q* → *R*(*p*(*q*)) is constant in a neighborhood of *q*_*_. Thus, the Jacobian *p*^*′*^(*q*_*_) spans the null-space of the Jacobian *J*(*p*_*_) of *R* at *p*_*_. A basis for the null-space of the Jacobian *J*(*p*_*_) can be computed, e.g., using the singular value decomposition (SVD) of *J*(*p*_*_). For any *d* ∈ ℝ ^*n*−*r*^, *p*(*q*_*_ + *d*) ≈ *p*(*q*_*_) + *p*^*′*^(*q*_*_)*d* and *R*(*p*(*q*_*_ + *d*)) ≈ *R*(*p*(*q*_*_)) + *J*(*p*(*q*_*_))*p*^*′*^(*q*_*_)*d* = *R*(*p*_*_) + *J*(*p*_*_)*p*^*′*^(*q*_*_)*d* = *R*(*p*_*_), because *p*^*′*^(*q*_*_) spans the null-space of the Jacobian *J*(*p*_*_). The linearization *p*(*q*_*_) + *p*^*′*^(*q*_*_)*d* is used to predict the next point *p*_**_ on the solution manifold. Given the predictor *p*(*q*_*_) + *p*^*′*^(*q*_*_)*d*, the new point *p*_**_ is computed as the solution of min 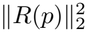 obtained using a nonlinear least-squares solver starting from the predictor. This is illustrated in Fig. 1D and summarized in Algorithm 2.

##### Algorithm 2

Continuation

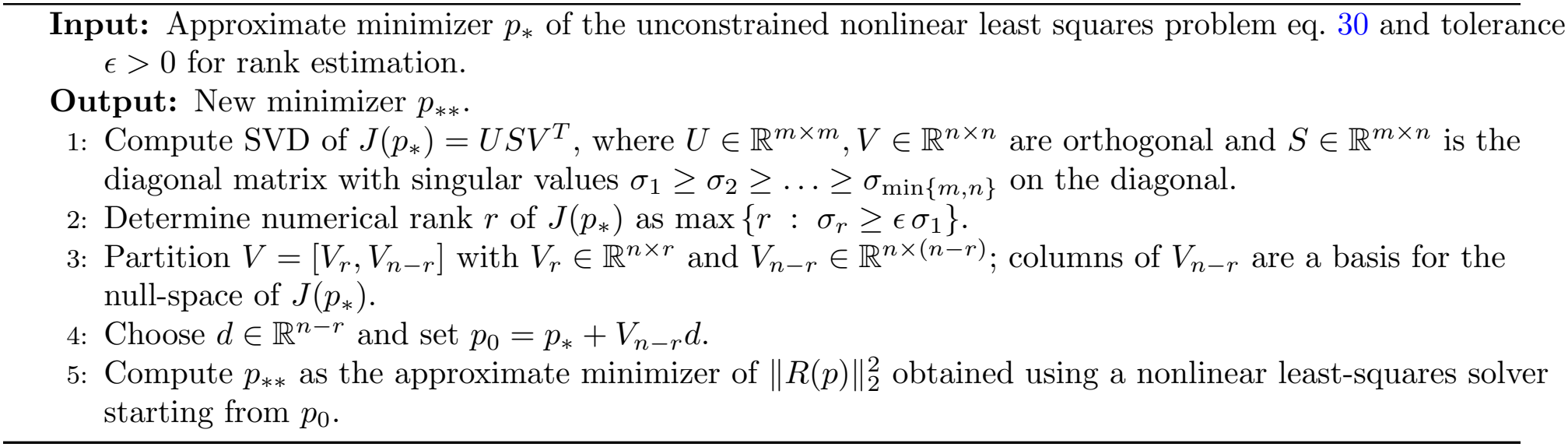

In our numerical experiments, we use a SciPy-based implementation of Algorithm 2, *ϵ* = 10^−12^, we set *d* to be a scalar *δ* times the (*n* − *r*)th unit vector so that *V*_*n*−*r*_*d* = *δv*_*n*_ ∈ ℝ ^*n*^ where *v*_*n*_ ∈ ℝ ^*n*^ is the right singular vector corresponding to the smallest singular value *σ*_*n*_, and we use <monospace>scipy.optimize.least_squares</monospace> with <monospace>method=‘trf’</monospace> as the nonlinear least-squares solver in Step 5.

For the LGMD H-current model, we performed 400 continuation steps with *δ* = 0.05, but more continuation steps would have been possible, in agreement with analytical results (sec. S.1). To trace the solution manifold for the A-current model, we performed 620 continuation steps with *δ* = 0.04. In this case, the continuation process could not be extended further. After adding tail current experiments, the extent of the solution manifold became negligible. For the HH model, the nonlinear least-squares solver used in Step 5 returned to the original local minimizer instead of converging to a new solution.

#### 4.2.5. Parameter confidence regions

When measurements are subject to measurement noise ***η*** that is normally distributed with mean zero and known symmetric positive definite covariance *C* ∈ ℝ^*m×m*^, i.e., ***η*** ∼ *N*_*m*_(0, *C*), then instead of eq. 30 we solve

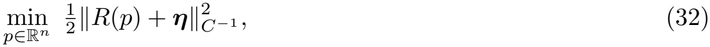

where 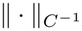 is the weighted Euclidean norm 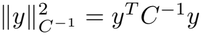. Let *p*^true^ ∈ ℝ^*n*^ be the solution of eq. 32 without noise, ***η*** 0. Note that for the cases handled in sec. 2, the covariance matrix *C* is proportional to the identity matrix. However, this may not always be the case, for example if the objective function includes membrane potential terms in addition to membrane current ones, see eq. 14.

Because of measurement noise, the solution **p** of eq. 32 is in general not equal to the true parameter *p*^true^, and confidence regions are subsets of ℝ^*n*^ around the solution of eq. 32 that contain the true parameter *p*^true^ with high probability See, e.g., [45, Section 7-10], [57], [58, Sec. 5.2]. Specifically, because of measurement noise, the solution **p** of eq. 32 is a random variable with *n* components. A (1 − *α*) *×* 100% confidence region for **p** is a subset CR_*α*_ ⊂ ℝ^*n*^ such that the probability of the true parameter *p*^true^ being in the confidence region CR_*α*_ is 1 − *α*, i.e.,

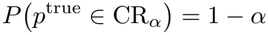

[57]. For linear least squares problems, i.e., *R*(*p*) = *Ap* + *b*, with normally distributed noise, the confidence region can be determined analytically, see, e.g., [57, Sec. 2]. A common way to compute approximate confidence regions in the nonlinear case is using linearizations and building on the linear case [57], [58, Sec. 5.2]. We summarize the approach next. In the linear case *R*(*p*) = *Ap* + *b*, all approximations below are exact. In the nonlinear case, the quality of approximations depends on the size of the ‘local curvature’ of *R*(*p*).

If the Jacobian of the residual function *R* : ℝ^*n*^ → ℝ^*m*^ at *p, J*(*p*) ∈ ℝ^*m×n*^, has full rank *n* for all *p*, then expanding the solution of eq. 32 with respect to ***η*** leads to the first order approximation

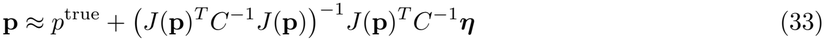

of the solution **p**. For a linear least squares problem, i.e., *R*(*p*) = *Ap* + *b*, eq. 33 is exact. If *p*_*η*_ is a computed realization of **p**, i.e., the solution of eq. 32 with a realization *η* of ***η***, then

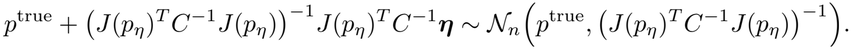

An approximate confidence region is computed by assuming that **p** is normally distributed with mean *p*^true^ and covariance (*J*(*p*_*η*_)^*T*^ *C*^−1^*J*(*p*_*η*_))^−1^, i.e., that

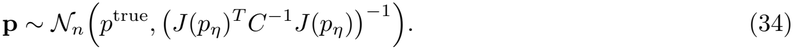

If eq. 34 holds, (*p*^true^ − **p**)^*T*^ *(J*(*p*_*η*_)^*T*^ *C*^−1^*J*(*p*_*η*_)) (*p*^true^ − **p**) can be written as the sum of *n* independent squared normally distributed random variables and therefore, has a chi-squared distribution with *n* degrees of freedom, 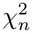. If *ρ*(*x*) is the probability density function of 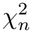, then for every 0 *< α <* 1, the quantile of the 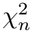 distribution at value 1 − *α* is *c*_1−*α*_ with 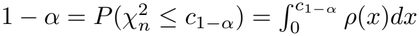. Thus, if eq. 34 holds, then *P* ((*p*^true^ − **p**)^*T*^ (*J*(*p*_*η*_)^*T*^ *C*^−1^*J*(*p*_*η*_)) (*p*^true^ − **p**) ≤ *c*_1−*α*_) = 1 − *α*. Therefore, the (approximate) (1 − *α*) *×* 100% confidence region is

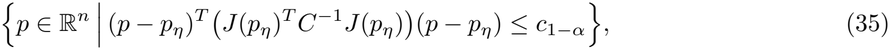

where, again, *p*_*η*_ is the parameter estimate, i.e., the solution of eq. 32 with a realization *η* of ***η***. The (approximate) confidence region eq. 35 is an ellipse, such as the ones shown in Fig. 3E and Fig. 5B. To obtain the (1 − *α*) *×* 100% confidence interval for a single parameter, the above is used restricted to a single parameter.

Note that we assume the symmetric positive definite covariance *C* ∈ ℝ^*m×m*^ to be known. In [57], [58, Sec. 5.2] it is assumed that *C* = *σ*^2^*I* with unknown *σ*^2^ *>* 0. In this case, eq. 35 is replaced by

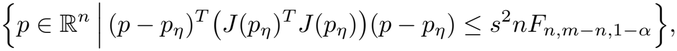

where 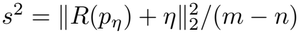 is the estimated residual variance, *p*_*η*_ is the solution of eq. 32 with *C* = *I* and realization *η* of ***η***, and *F*_*n,m*−*n*,1−*α*_ is the 1 − *α* quantile of the *F*_*n,m*−*n*_ distribution.

#### 4.2.6. Hierarchical clustering of parameter correlations

We computed parameter correlations from the covariance matrix at the local minimum *p*, where the covariance matrix was approximated using (*J*^*T*^ (*p*)*J*(*p*))^−1^. The correlation coefficient between parameters *i* and *j* is denoted by *ρ*_*i,j*_. Pairwise parameter distances were then defined as *d*_*i,j*_ = 1 − |*ρ*_*i,j*_| . We performed hierarchical clustering on the resulting distance matrix using the average linkage method implemented in <monospace>scipy.cluster.hierarchy</monospace>.

## Supporting information

### S.1 LGMD H-current model rank deficiency under voltage clamp

Although the number of measurements, *n*_*V*_ *m*_*V*_, is much larger than the number of parameters to be estimated, 5 + 2*n*_*V*_ or 4 + 2*n*_*V*_ if 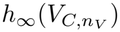 is fixed, the Jacobian associated with the nonlinear least squares problem eq. 8 is rank deficient. This rank deficiency implies that the parameters cannot be uniquely identified from eq. 8. The following result (proven in [21]) states that for the voltage clamp experiments and a wide range of biophysically relevant parameters, the Jacobian has a rank deficiency of three before fixing 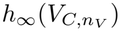. After fixing and excluding 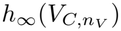 from the estimation problem, the rank deficiency is reduced to two.

#### Assumption S.1

*The command voltages satisfy V*_*C,i*_≠ *V*_*C,j*_ *for i*≠ *j, and the parameters eq. 6 satisfy* 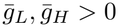, *V*_*C,i*_ *E*_*H*_, *h*_∞_(*V*_*C,i*_)≠ *h*_∞_(*V*_*C*,hold_), *and τ*_*h*_(*V*_*C,i*_) *>* 0, *i* = 1, …, *n*_*V*_ .

#### Theorem S.2

*Let Assumption S.1 hold, n*_*V*_ *>* 1, *and m*_*V*_ *>* 2.

- *The Jacobian* 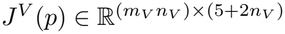 *corresponding to eq. 8 has rank* 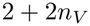
- *The Jacobian* 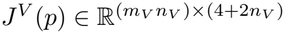 *corresponding to eq. 8 with* 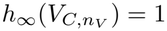 *and parameters eq. 7 has rank* 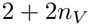.

Figure 1C shows the normalized singular values for *n*_*V*_ = 5. If the parameters eq. 6 are used, the smallest three normalized singular values (black crosses) are below *ϵ* = 10^−12^. This matches the theory in Theorem S.2i, which states that the rank deficiency of *J*^*V*^ (*p*) is three. If the parameters eq. 7 are used, the smallest two normalized singular values (green dots) are below *ϵ* = 10^−12^. This matches the theory in Theorem S.2ii, which states that the rank deficiency of *J*^*V*^ (*p*) is two.

### S.2 Systematic errors from incorrect parameter constraints

Note that assuming 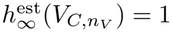 when the true parameter satisfies 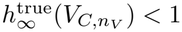 introduces a systematic error. To see this, assume that the true parameters solve eq. 8. However, if we fix 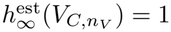 then solving eq. 8 leads to

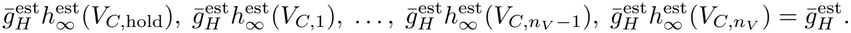

Comparing these estimates with

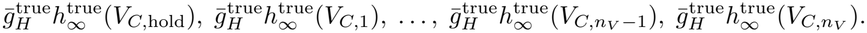

shows that the estimated parameters are scaled versions of the true parameters

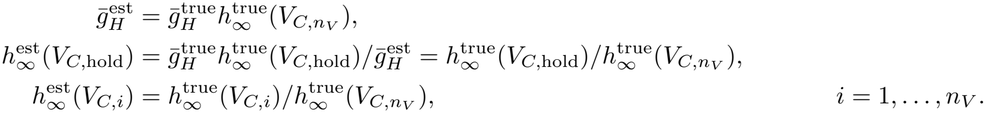

### S.3 LGMD H-current model rank deficiency after tail experiments

Adding tail current experiments increases the number of parameters to be identified, but reduces the rank deficiency of the Jacobian associated with the nonlinear least squares problem eq. 11. The following result (proven in [21]) states that adding tail current experiments reduces the rank deficiency of the Jacobian to two and to one if 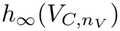 is fixed and excluded from the parameters.

#### Assumption S.3

*Assumption S.1 holds. Furthermore, the command voltages satisfy V*_*T,i*_≠ *V*_*T,j*_ *for i*≠ *j, and the parameters satisfy V*_*T,i*_≠ *E*_*H*_, *h*_∞_(*V*_*T,i*_)≠ *h*_∞_(*V*_*T*,hold_), *h*_∞_(*V*_*C*,hold_)≠ *h*_∞_(*V*_*T*,hold_), *and τ*_*h*_(*V*_*T,i*_) *>* 0, *i* = 1, …, *n*_*T*_ .

#### Theorem S.4

*Let Assumption S.3 hold, n*_*V*_ *>* 1, *m*_*V*_ *>* 2, *n*_*T*_ *>* 1, *and m*_*T*_ *>* 2.

i. *The Jacobian* 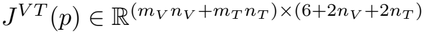 *has rank* 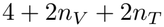
ii. *The Jacobian* 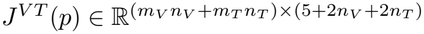 *corresponding to eq. 11 with* 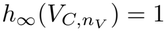 *and parameters eq. 10 has rank* 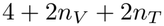.

As before, the rank of the Jacobian *J*^*VT*^ (*p*) can be seen from its singular values. Figure 1C shows the normalized singular values for *n*_*V*_ = 5 and *n*_*T*_ = 4 using parameters eq. 10. The smallest normalized singular value (blue squares) is below *ϵ* = 10^−12^. This matches the theory in Theorem S.4ii, which states that the rank deficiency of *J*^*VT*^ (*p*) is one.

Adding a passive current clamp experiment further removes the rank deficiency, resulting in a full-rank Jacobian.

### S.4. LGMD H-current model rank after current clamp experiments

Adding passive current clamp experiments does not increase the number of parameters to be identified, but further reduces the rank deficiency of the Jacobian associated with the nonlinear least squares problem eq. 14. The following result (proven in [21]) states that adding current clamp experiments reduces the rank deficiency of the Jacobian to one, and that the Jacobian has full column rank if 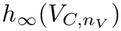 is fixed and excluded from the parameters.

#### Assumption S.5

*Assumption S.3 holds. Furthermore, let n*_*V*_ *>* 1, *m*_*V*_ *>* 2, *and let either (a) n*_*C*_ ≥ 2 *and I*_*C,i*_≠ *I*_*C,j*_ *for i*≠ *j, or (b) m*_*C*_ ≥ 2 *and* 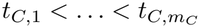.

#### Theorem S.6

*Let Assumption S.5 hold, n*_*T*_ *>* 1, *and m*_*T*_ *>* 2.

i. *The Jacobian* 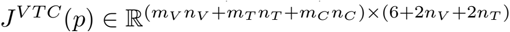 *corresponding to eq. 14 with parameters eq. 9 has rank* 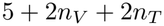.
ii. *The Jacobian* 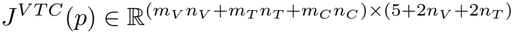 *corresponding to eq. 14 with* 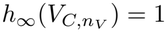 *and parameters eq. 10 has full column rank* 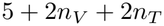.

As before, the rank of the Jacobian *J*^*VTC*^(*p*) can be seen from its singular values. Figure 1C shows the normalized singular values for *n*_*V*_ = 5, *n*_*T*_ = 4, and *n*_*C*_ = 3 using parameters eq. 10. None of the normalized singular values (red trace) is below *ϵ* = 10^−12^. This matches the theory in Theorem S.6ii, which states that *J*^*VTC*^(*p*) has full column rank.

### S.5. CA1 H-current model rank deficiency

Surprisingly, if the leak channel is not present in the model, the rank result for the Jacobian of the least squares problem corresponding to eq. 8 with parameters eq. 16 or with 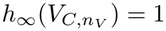 fixed is substantially different than those reported in Theorem S.2. Recall that the least squares problem corresponding to eq. 8 with parameters eq. 16 has *n*_*V*_ *m*_*V*_ measurements and 3 + 2*n*_*V*_ parameters, or, with 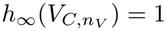 fixed, 2 + 2*n*_*V*_ parameters. The rank of the Jacobian now depends on whether *h*_∞_(*V*_*C*,hold_) = 0 or *h*_∞_(*V*_*C*,hold_) *>* 0. The following result, proven in [21], states that for the voltage clamp experiments and a wide range of biophysically relevant parameters, the Jacobian has a rank deficiency of two if *h*_∞_(*V*_*C*,hold_) = 0 but a rank deficiency of one if *h*_∞_(*V*_*C*,hold_) *>* 0. If 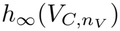 is fixed and removed from the parameters, the Jacobian has a rank deficiency of one if *h*_∞_(*V*_*C*,hold_) = 0 but has full column rank if *h*_∞_(*V*_*C*,hold_) *>* 0.

#### Theorem S.7

*Let the command voltages and the parameters of eq. 16 satisfy Assumption S.1, and n*_*V*_ *>* 1, *m*_*V*_ *>* 2.

*i.a. If h*_∞_(*V*_*C*,hold_) = 0, *then the Jacobian* 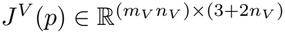 *corresponding to eq. 8 has rank* 1 + 2*n*_*V*_ .

*i.a. If h*_∞_(*V*_*C*,hold_) *>* 0, *then the Jacobian* 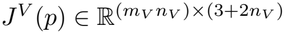 *corresponding to eq. 8 has rank* 2 + 2*n*_*V*_ .

*ii.b. If* 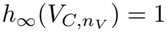 *is fixed and h*_∞_(*V*_*C*,hold_) = 0, *then the Jacobian* 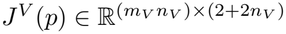 *corresponding to eq. 8 with parameters eq. 16 (with* 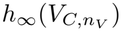 *removed) has rank* 1 + 2*n*_*V*_ .

*ii.b. If* 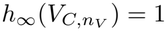 *is fixed and h*_∞_(*V*_*C*,hold_) *>* 0, *then the Jacobian* 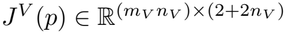 *corresponding to eq. 8 with parameters eq. 16 (with* 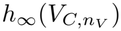 *removed) has full rank* 2 + 2*n*_*V*_ .

Again, the rank of the Jacobian *J*^*V*^ (*p*) can be seen from its singular values. Figure 2C shows the normalized singular values for *n*_*V*_ = 7 and for the case where 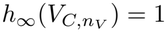 is fixed. In the noise-free case *h*_∞_(*V*_*C*,hold_) = 0, and the smallest normalized singular value is below *ϵ* = 10^−12^ (green dots, Fig. 2C). This matches the theory in Theorem S.7iia, which states that the rank deficiency of *J*^*V*^ (*p*) is one. If the measurements are subject to noise, then this noise impacts the computed parameter, which results in *h*_∞_(*V*_*C*,hold_) *>* 0. In this case, all normalized singular values are above 10^−9^ (red dots, Fig. 2C). This matches the theory in Theorem S.7iib, which states that the Jacobian has full rank. The rank of the Jacobian depends on whether *h*_∞_(*V*_*C*,hold_) = 0 or *h*_∞_(*V*_*C*,hold_) *>* 0, and the observed change in rank changes because measurement noise results in an estimated *h*_∞_(*V*_*C*,hold_) *>* 0.

### S.6 CA1 H-current model rank after tail current experiments

Fitting the tail portion of the voltage clamp experiment, i.e., the current obtained when the membrane potential is clamped back from each command voltage *V*_*C,i*_ to the holding potential *V*_*C*,hold_, adds *n*_*V*_ *m*_*T*_ measurements but only one parameter, *τ*_*h*_(*V*_*C*,hold_), to the estimation problem. Thus, the least squares problem corresponding to eq. 11 with model eq. 15 has 4 + 2*n*_*V*_ parameters, given by eq. 16 plus *τ*_*h*_(*V*_*C*,hold_).

With 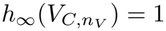 fixed, it has 3 + 2*n*_*V*_ parameters, given by eq. 17. The following result (proven in [21]) states that adding tail current experiments reduces the rank deficiency of the Jacobian to one, and that the Jacobian has full column rank if 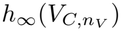 is fixed and excluded from the parameters. In contrast to the voltage clamp experiment alone (sec. S.5), this result holds whether *h*_∞_(*V*_*C*,hold_) = 0 or *h*_∞_(*V*_*C*,hold_) *>* 0.

#### Theorem S.8

*Let the command voltages and parameters eq. 16 satisfy Assumption S.1, and τ*_*h*_(*V*_*C*,hold_) *>* 0, *m*_*V*_ *>* 2, *m*_*T*_ *>* 2.

i. *The Jacobian* 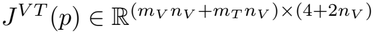 *corresponding to the parameters eq. 16 and τ*_*h*_(*V*_*C*,hold_) *has rank* 3 + 2*n*_*V*_ .
ii. *The Jacobian* 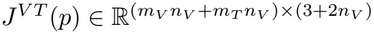 *corresponding to* 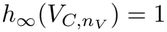 *and parameters eq. 17 has full column rank* 3 + 2*n*_*V*_ .

As before, the rank of the Jacobian *J*^*VT*^ (*p*) can be seen from its singular values. Figure 2C shows the normalized singular values for *n*_*V*_ = 7 using parameters eq. 17. None of the normalized singular values (blue dots) is below *ϵ* = 10^−12^. This matches the theory in Theorem S.8ii, which states that *J*^*VT*^ (*p*) has full column rank.

### S.7. A-current model rank under voltage clamp

In the A-current model eq. 18, the maximal conductance 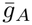 multiplies the activation and the inactivation variables. This leads to a model-based multiplicative parameter degeneracy. In contrast to the LGMD H-current model (sec. S.1), this model has no additional degeneracy involving the reversal potentials. The following result (proven in [21]) states that the resulting rank deficiency of the Jacobian is two, and that the Jacobian has full column rank if *h*_∞_(*V*_*C*,hold_) and 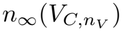 are fixed and excluded from the parameters

#### Assumption S.9

*Assume that n*_*V*_ *>* 1, *the sample times* 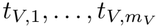 *are distinct with m*_*V*_ ≥ 15, *V*_*C,i*_≠ *V*_*C,j*_ *for i*≠ *j, and the parameters eq. 19 satisfy* 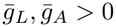, *V*_*C,i*_≠ *E*_*A*_, *h*_∞_(*V*_*C*,hold_)≠ 0, *n*_∞_(*V*_*C*,hold_)≠ 0, *h*_∞_(*V*_*C,i*_)≠ *h*_∞_(*V*_*C*,hold_), *n*_∞_(*V*_*C,i*_)≠ *n*_∞_(*V*_*C*,hold_), *τ*_*h*_(*V*_*C,i*_) *>* 0, *τ*_*n*_(*V*_*C,i*_) *>* 0, *τ*_*h*_(*V*_*C,i*_)≠ *τ*_*n*_(*V*_*C,i*_), *h*_∞_(*V*_*C,i*_)≠ 0 *or n*_∞_(*V*_*C,i*_)

#### Theorem S.10

*Assumption S.9 holds*.

1. *The Jacobian* 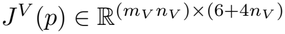 *corresponding to eq. 8 with model eq. 18 and parameters eq. 19 has rank* 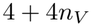.
2. *The Jacobian* 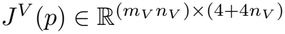 *corresponding to eq. 8 with model eq. 18, h*_∞_(*V*_*C*,hold_) = 1, 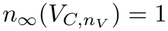, *and parameters eq. 20 has full column rank* 4 + 4*n*_*V*_ .

As before, the rank of the Jacobian *J*^*V*^ (*p*) can be seen from its singular values. Figure 3B shows the normalized singular values for *n*_*V*_ = 5 using parameters eq. 20. None of the normalized singular values (blue dots) is below *ϵ* = 10^−12^. This matches the theory in Theorem S.10ii, which states that *J*^*V*^ (*p*) has full column rank.

### S.8 Rate functions of the Hodgkin Huxley model

The general form of the rate functions used in [1] is as follows:

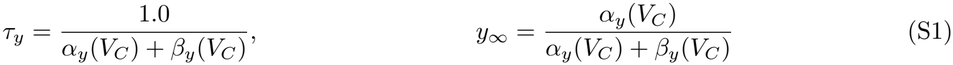

The rate functions of the sodium current are

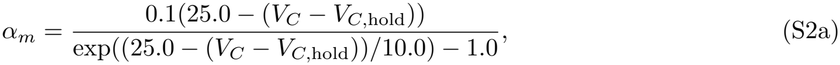

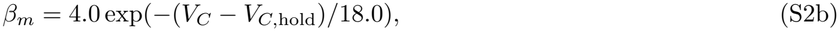

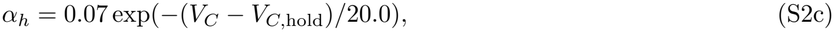

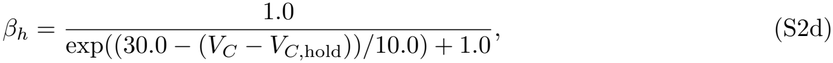

and those of the potassium current are

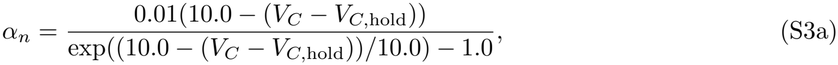

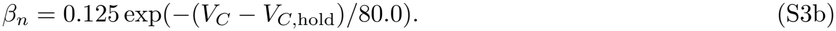

### S.9 Hodgkin-Huxley sodium and potassium current model ranks

The sodium current model eq. 22 has the same functional form as the A-current model eq. 18, with *n* and *h* replaced by *m* and *h*, respectively. Therefore, Theorem S.10 applies [21]: the Jacobian corresponding to eq. 8 with model eq. 22 and parameters eq. 23 has rank 4 + 4*n*_*V*_, and the Jacobian with parameters eq. 24, i.e., after fixing *h*_∞_(*V*_*C*,hold_) = 0.6 and 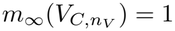, has full column rank. Figure 4D shows the normalized singular values for *n*_*V*_ = 7 using parameters eq. 24. None of the normalized singular values (blue dots) is below *ϵ* = 10^−12^, in agreement with this result.

For the potassium current model, the following result (proven in [21]) states that the Jacobian has a rank deficiency of one. After fixing 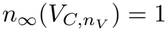 and excluding it from the estimation problem, the Jacobian is full rank.

#### Assumption S.11

*Assume that n*_*V*_ *>* 1, *the sample times* 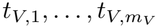 *are distinct with m*_*V*_ ≥ 9, *V*_*C,i*_≠ *V*_*C,j*_ *for i*≠ *j, and the parameters eq. 26 satisfy* 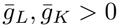, *V*_*C,i*_ *n*_∞_(*V*_*C,i*_)≠ *n*_∞_(*V*_*C*,hold_), *τ*_*n*_(*V*_*C,i*_) *>* 0, *i* = 1, …, *n*_*V*_ .

#### Theorem S.12

*Assumption S.11 holds*.

i. *The Jacobian* 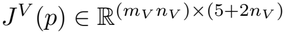 *corresponding to eq. 8 with model eq. 25 and parameters eq. 26 has rank* 4 + 2*n*_*V*_ .
ii. *The Jacobian* 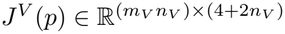 *corresponding to eq. 8 with model eq. 25*, 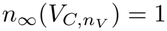 *and parameters eq. 27 has full column rank* 4 + 2*n*_*V*_ .

Figure 4E shows the normalized singular values for *n*_*V*_ = 7 using parameters eq. 27. None of the normalized singular values (blue dots) is below *ϵ* = 10^−12^. This matches the theory in Theorem S.12ii, which states that *J*^*V*^ (*p*) has full column rank. If the fourth-order activation is replaced by a first-order one, the model has the same form as the LGMD H-current model eq. 5, and the smallest two normalized singular values (red dots) are below *ϵ* = 10^−12^. This matches the theory in Theorem S.2ii, which states that the rank deficiency of *J*^*V*^ (*p*) is two.

### S.10 Figures

Supplementary figures provide additional results and analyses that complement the findings presented in Section 2.

**Fig S1.**
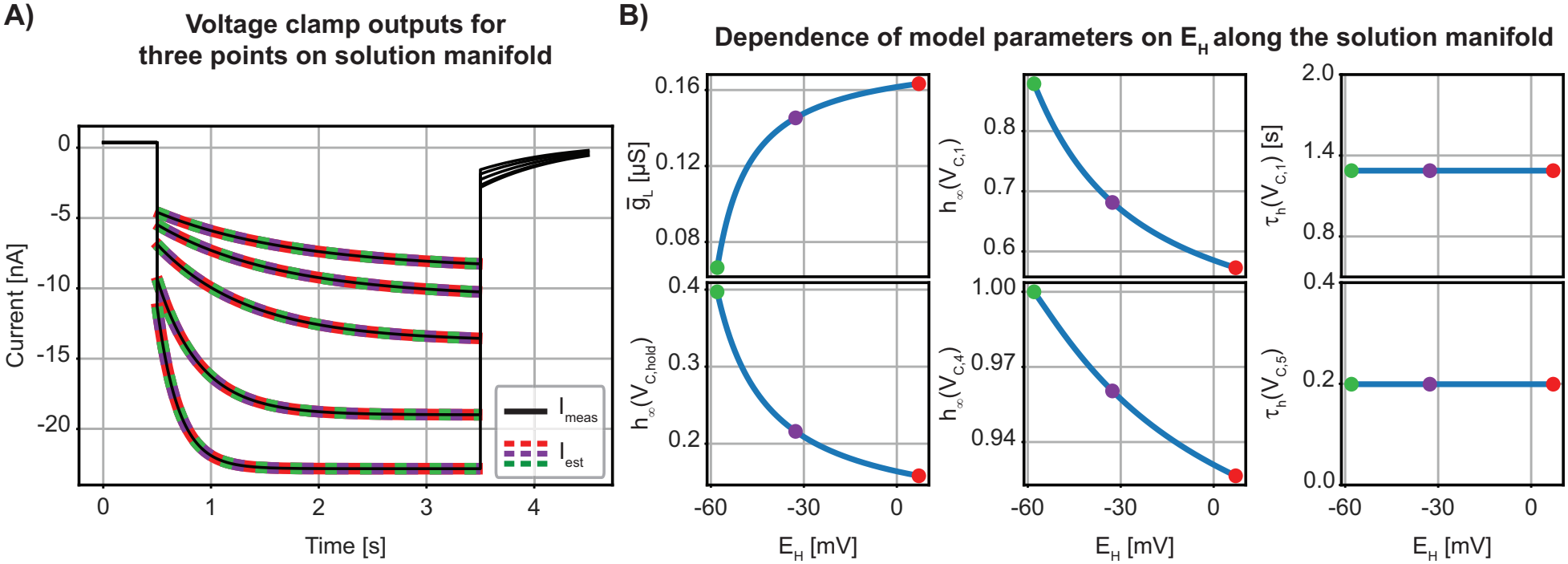
Additional examples illustrating parameter degeneracy in the LGMD H-current model. **(A)** Voltage clamp outputs for the three colored dots highlighted in Fig. 1E. Black traces show the measured currents used for parameter estimation, and dashed colored lines show the corresponding model outputs. Despite substantial differences in parameter values, the three model outputs are nearly identical. **(B)** Dependence of selected parameters on *E*_*H*_ for fixed *E*_*L*_ along the solution manifold.

**Fig S2.**
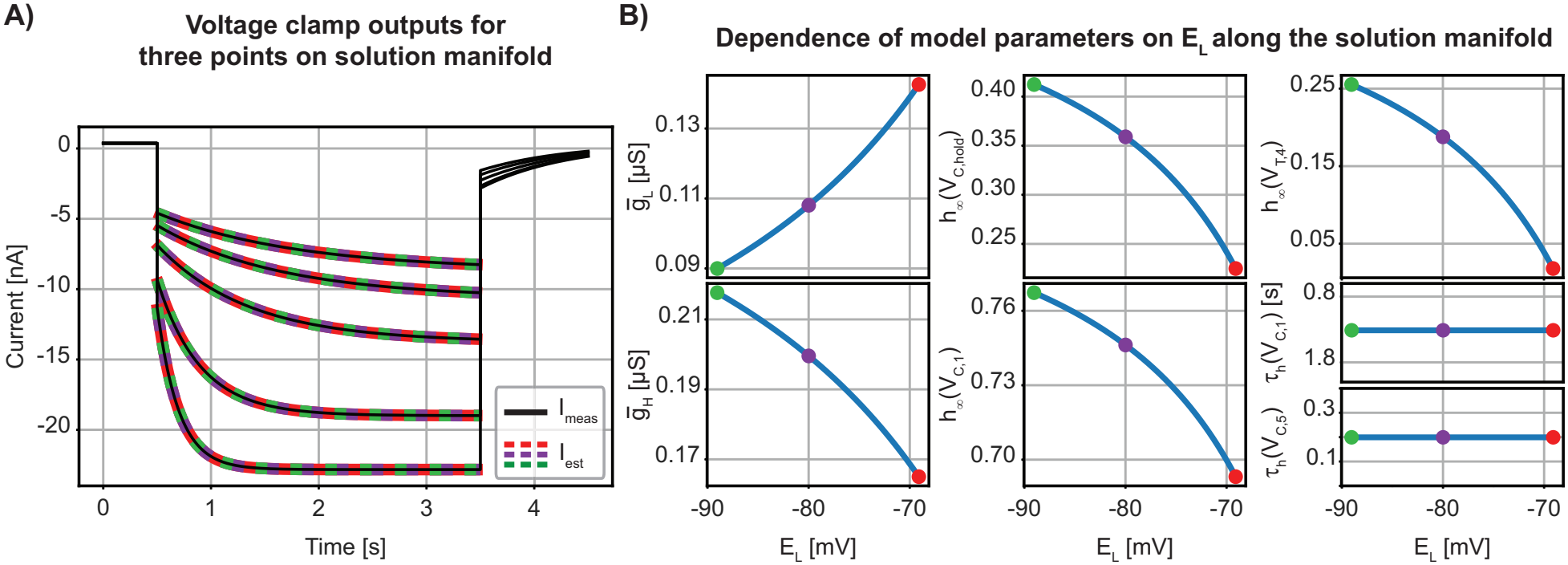
Illustrating parameter degeneracy in the LGMD H-current model after fixing *E*_*H*_ using the tail current experiment. **(A)** Voltage clamp outputs for the three colored dots highlighted in (B). Black traces show the measured currents used for parameter estimation, and dashed colored lines show the corresponding model outputs. Despite significant differences in parameter values, the three model outputs are nearly identical. (B) Dependence of selected parameters on *E*_*L*_ along the solution manifold with *E*_*H*_ fixed at −35 [mV] by adding the tail current experiment.

**Fig S3.**
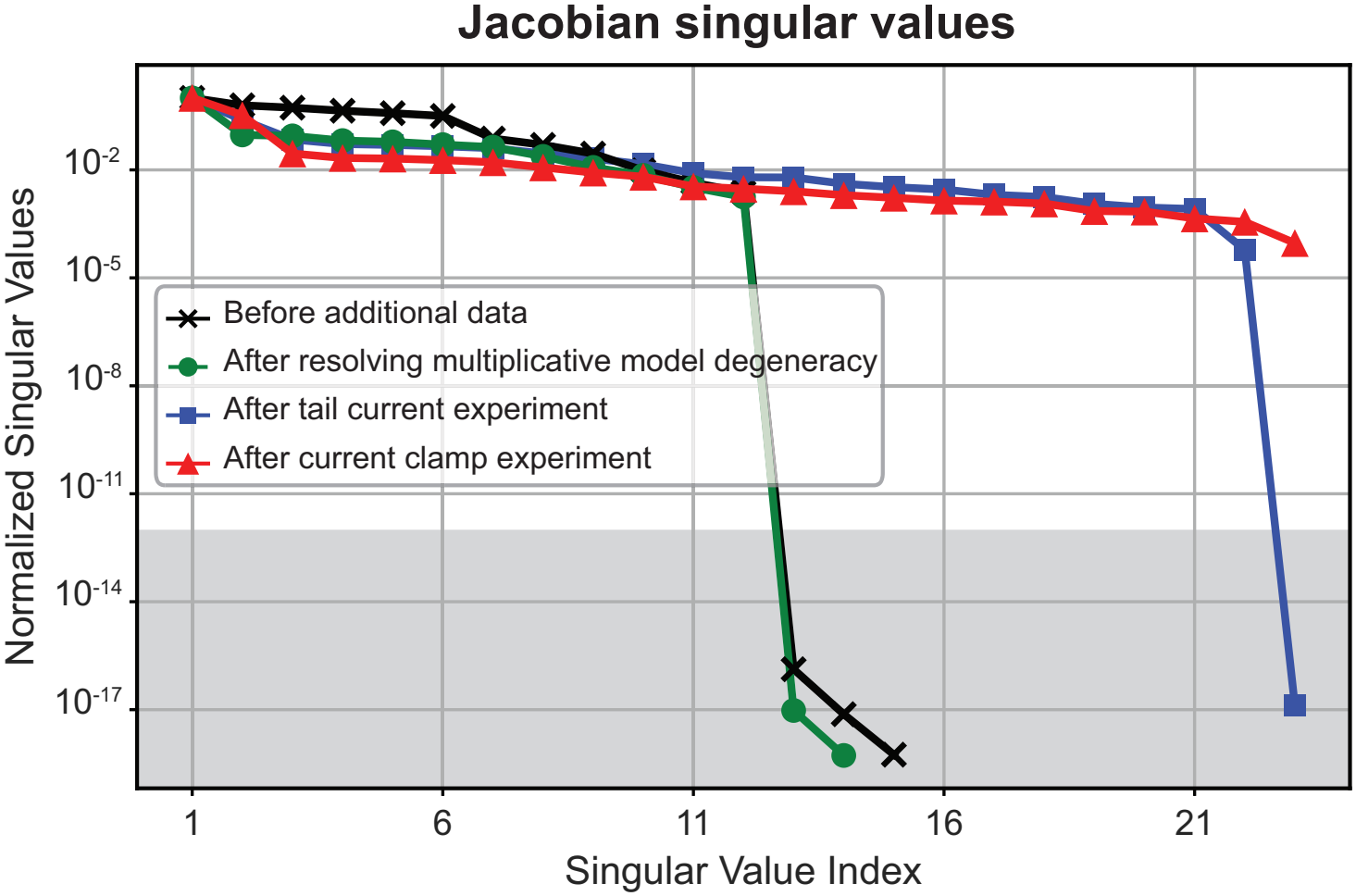
Normalized Jacobian singular values at different stages of the model determination process for noisy data. Singular values that fall within the gray region, below the threshold of 10^−12^, are considered to be equal to zero. The addition of noise to this model does not change the results qualitatively compared with the noise-free case shown in Fig. 1C.

**Fig S4.**
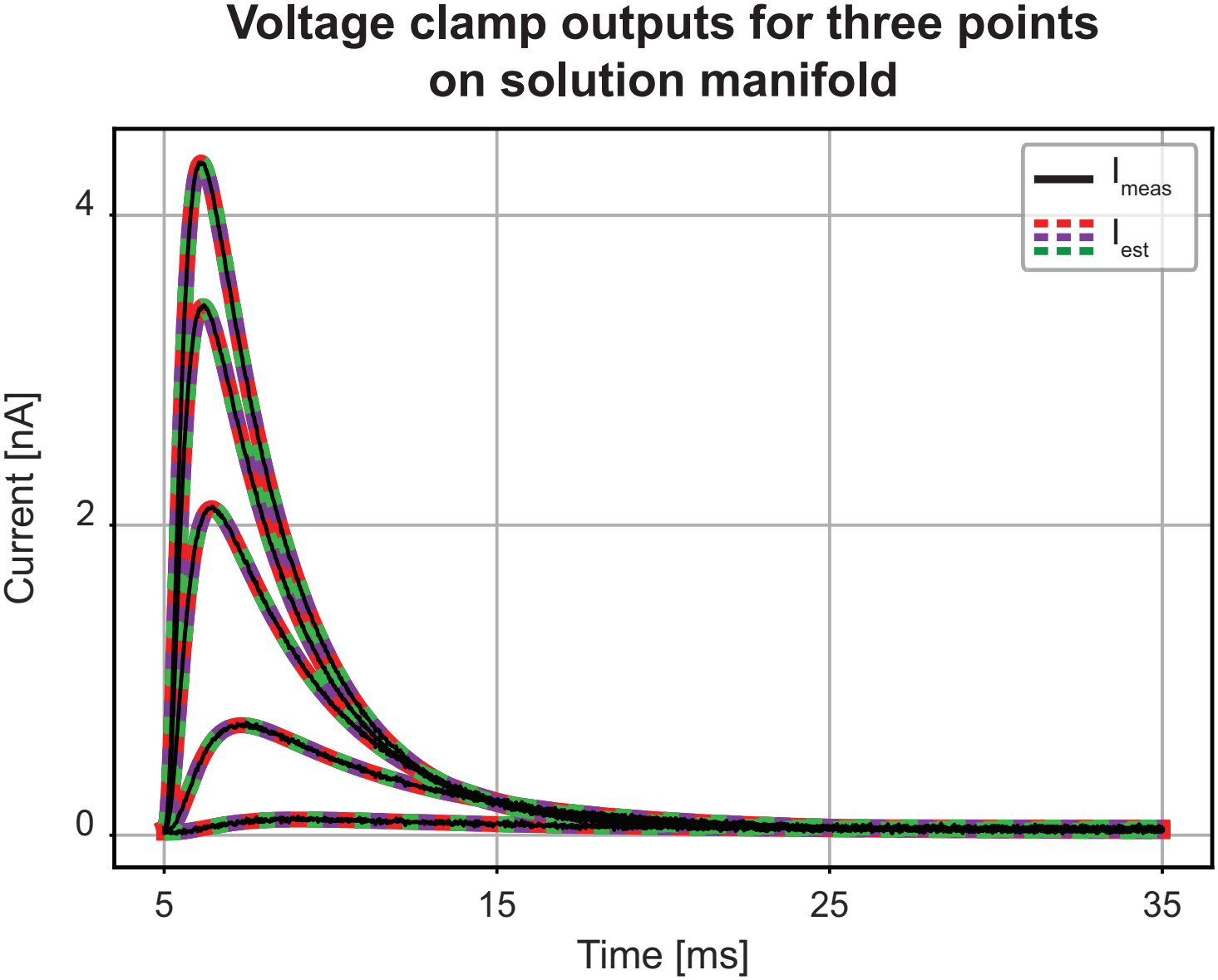
Voltage clamp outputs corresponding to the three parameter sets marked on the solution manifold in Fig. 3C. Despite differences in parameter values, all three parameter sets produce similar voltage clamp outputs, illustrating the model’s parameter degeneracy.

**Fig S5.**
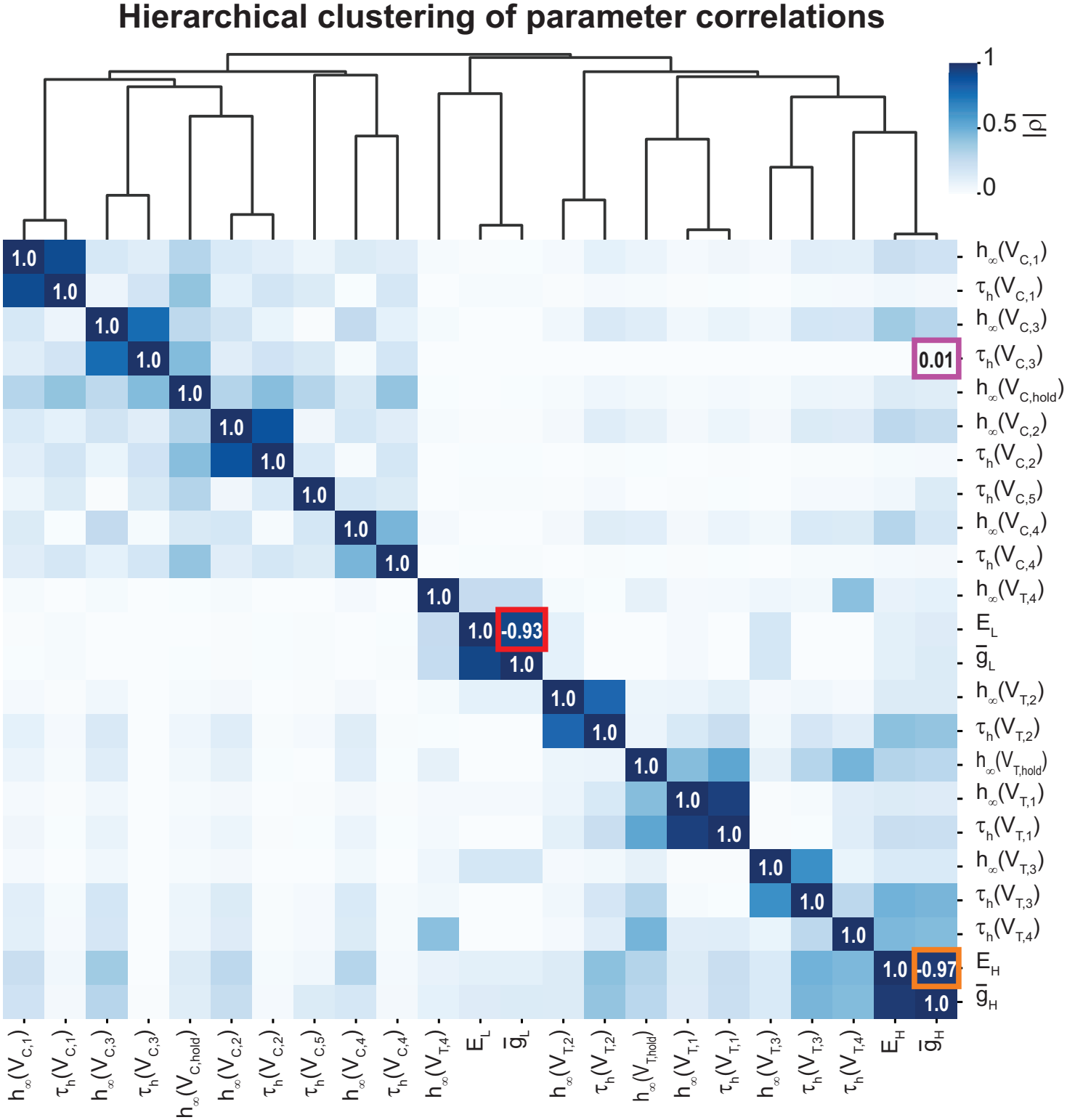
Hierarchical clustering of the correlation coefficient matrix for the noisy LGMD H-current model. The clustering analysis reveals relationships among parameters and identifies groups of highly correlated parameters. The orange and purple boxes mark the two parameter pairs whose projected 95% confidence regions are shown in Fig. S6. Consistent with the CA1 H-current model (see Fig. 5), the reversal potential and maximal conductance of an ionic current are highly correlated, as indicated by the red and orange boxes.

**Fig S6.**
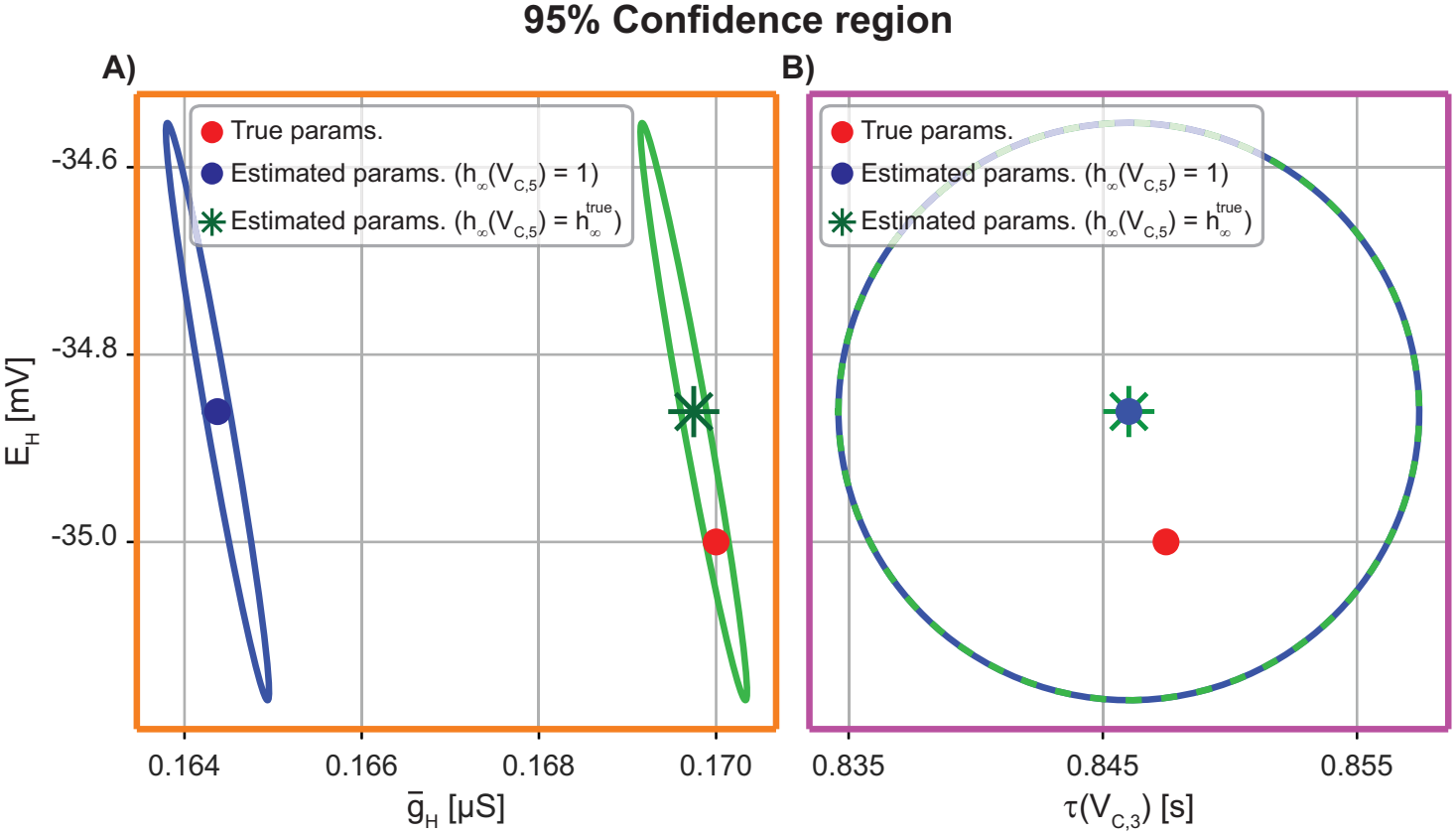
95% confidence regions for selected parameter pairs in the noisy LGMD H-current model. **(A)** Projected 95% confidence region for the strongly correlated parameters *E*_*H*_ and 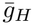, corresponding to the orange box in Fig. S5. The elongated and tilted ellipse reveals a direction of uncertainty in parameter space that is not apparent from the individual confidence intervals alone. The blue region was obtained by fixing *h*_∞_(*V*_*C*,5_) to one, to avoid multiplicative model-based degeneracy, while the green region was obtained by fixing it to its true value. Assuming *h*_∞_(*V*_*C*,5_) equals one scales 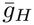 as these two parameters appear as products in our model. This scaling causes the true value to fall outside the blue confidence region. Projected 95% confidence regions for the weakly correlated parameters, corresponding to the purple box in Fig. S5. The nearly circular confidence region reflects nearly independent parameter uncertainties. Since the selected parameters do not appear as products in the model, fixing *h*_∞_(*V*_*C*,5_) does not affect their estimation. Thus, the dashed blue and green confidence regions overlap.

**Fig S7.**
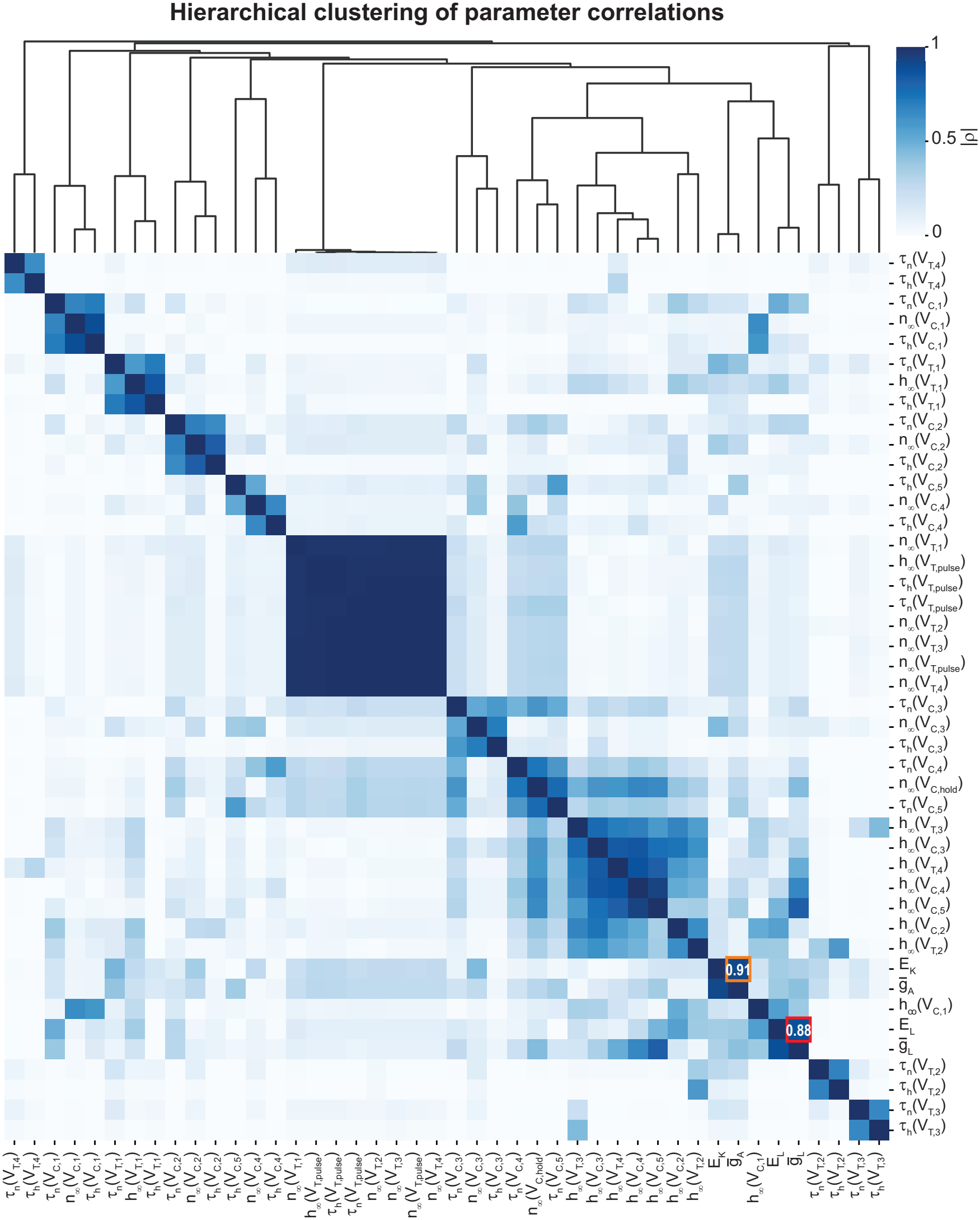
Hierarchical clustering of the correlation coefficient matrix for the A-current model. The clustering analysis reveals relationships among parameters and identifies groups of highly correlated parameters. Consistent with the CA1 H-current model (see Fig. 5), the reversal potential and maximal conductance of an ionic current are highly correlated, as indicated by the red and orange boxes.

**Fig S8.**
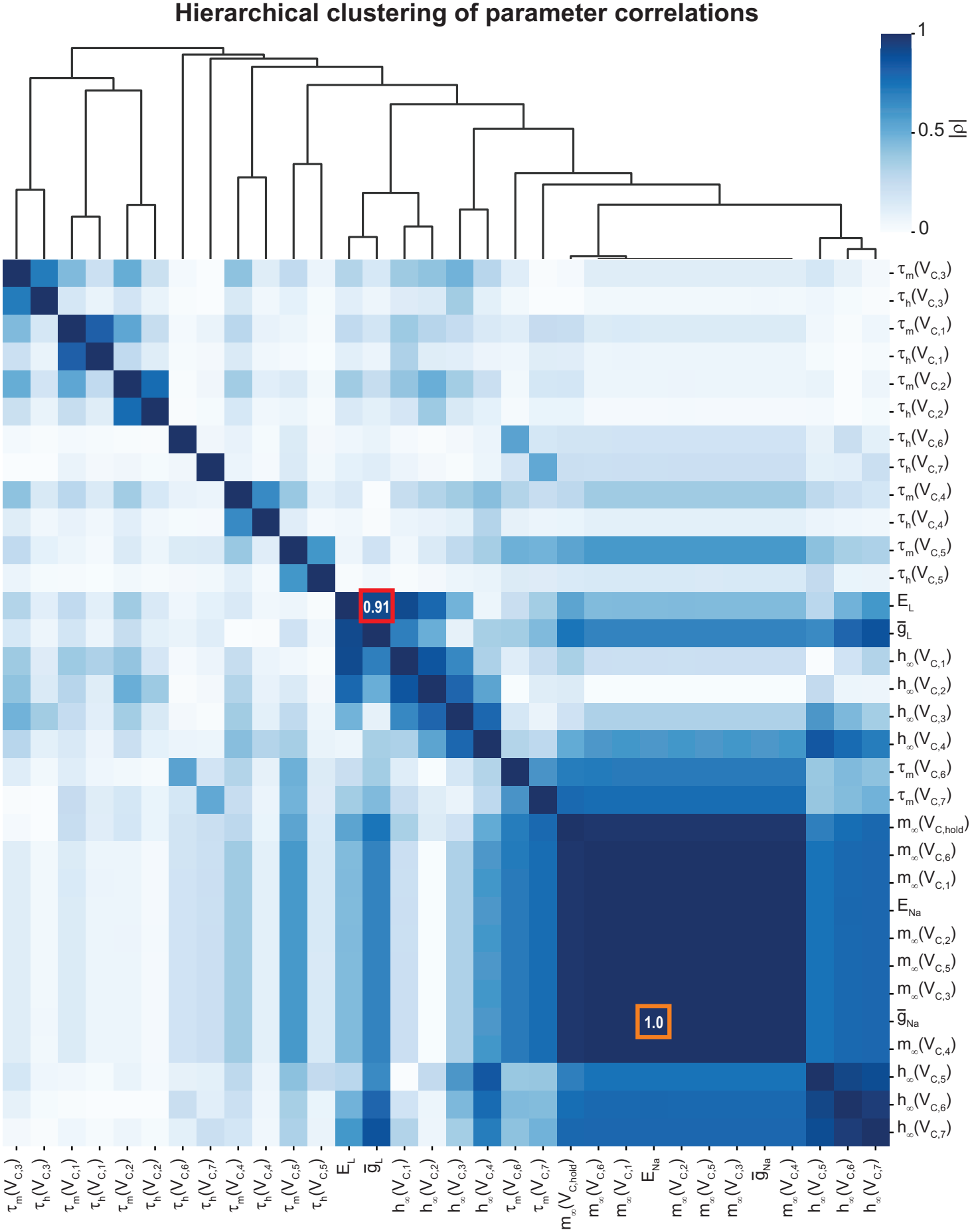
Hierarchical clustering of the correlation coefficient matrix for the sodium current (*I*_*Na*_) model. The clustering analysis reveals relationships among parameters and identifies groups of highly correlated parameters. Consistent with the CA1 H-current model (see Fig. 5), the reversal potential and maximal conductance of an ionic current are highly correlated, as indicated by the red and orange boxes.

**Fig S9.**
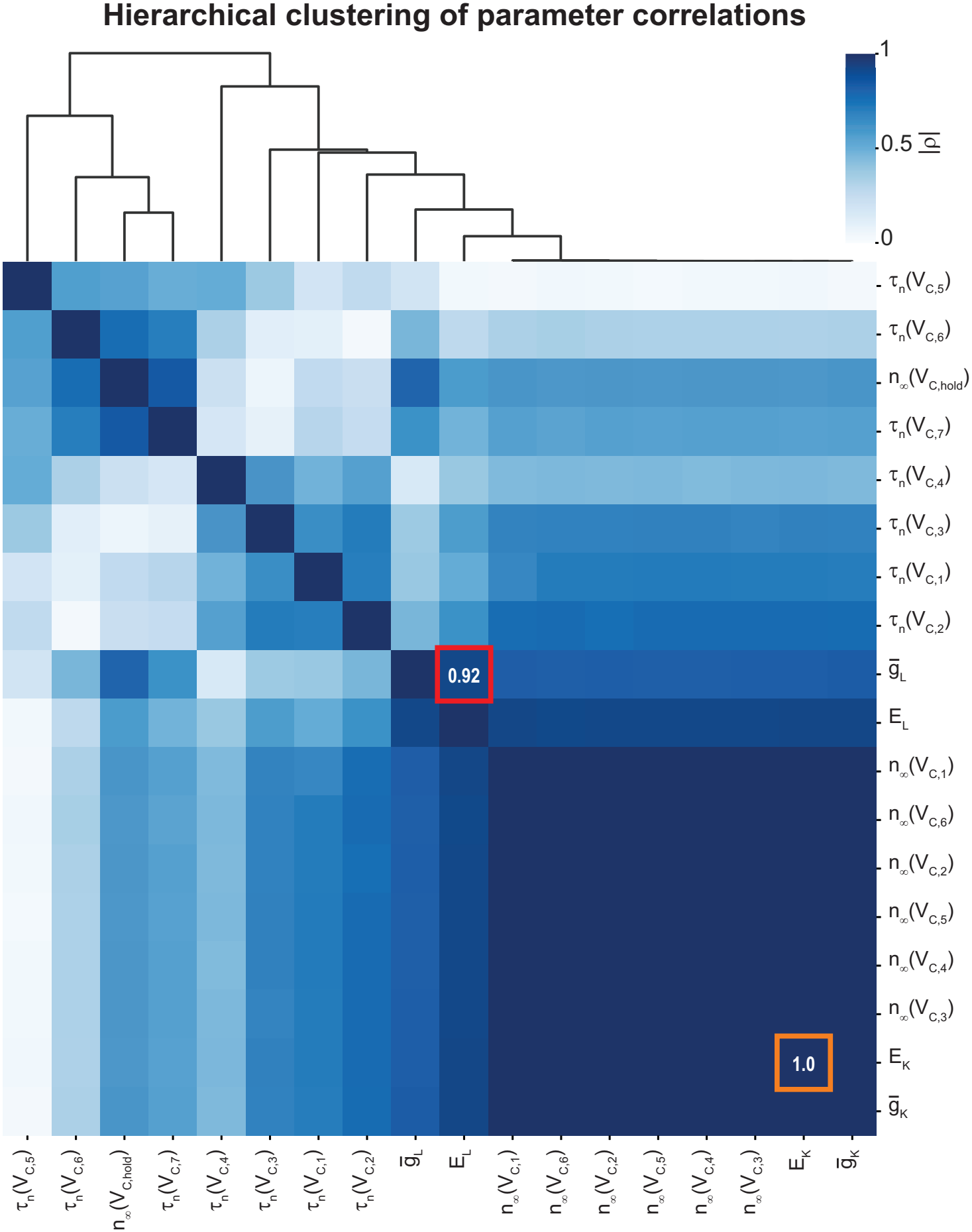
Hierarchical clustering of the correlation coefficient matrix for the potassium current (*I*_*K*_) model. The clustering analysis reveals relationships among parameters and identifies groups of highly correlated parameters. Consistent with the CA1 H-current model (see Fig. 5), the reversal potential and maximal conductance of an ionic current are highly correlated, as indicated by the red and orange boxes.

**Fig S10.**
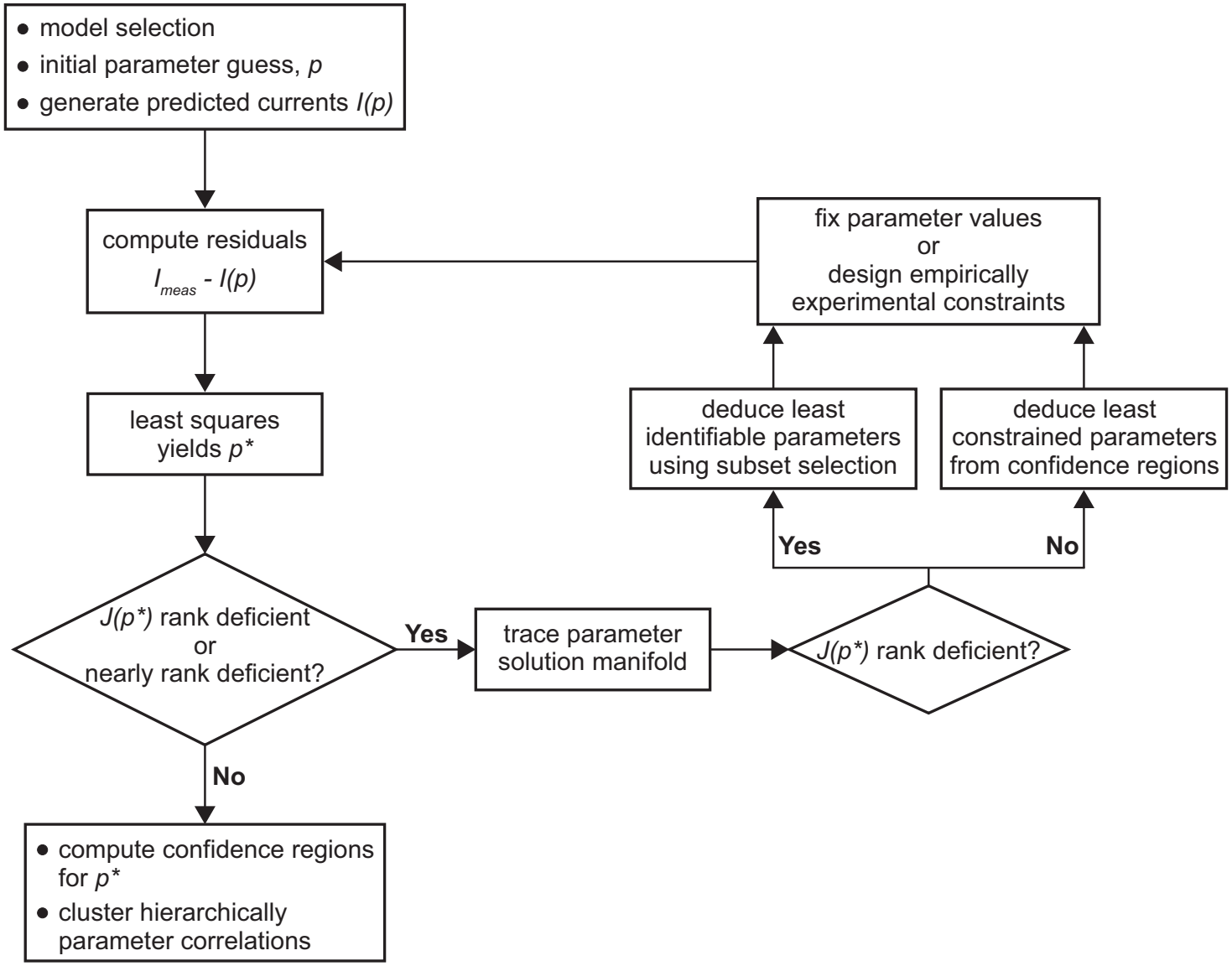
A toolkit identifying parameter degeneracy in HH-type current models. Organogram summarizing the steps developed and applied for the identification of model parameter degeneracy and solution manifolds of HH-type models in sec. 2.

### S.11. Tables

Specification of the true parameter values used to generate the synthetic data for the models described in Section 4 is provided in Tables S1, S3, S4, and S5. An example of the parameter estimation results is provided in Table S2.

**Table S1.**
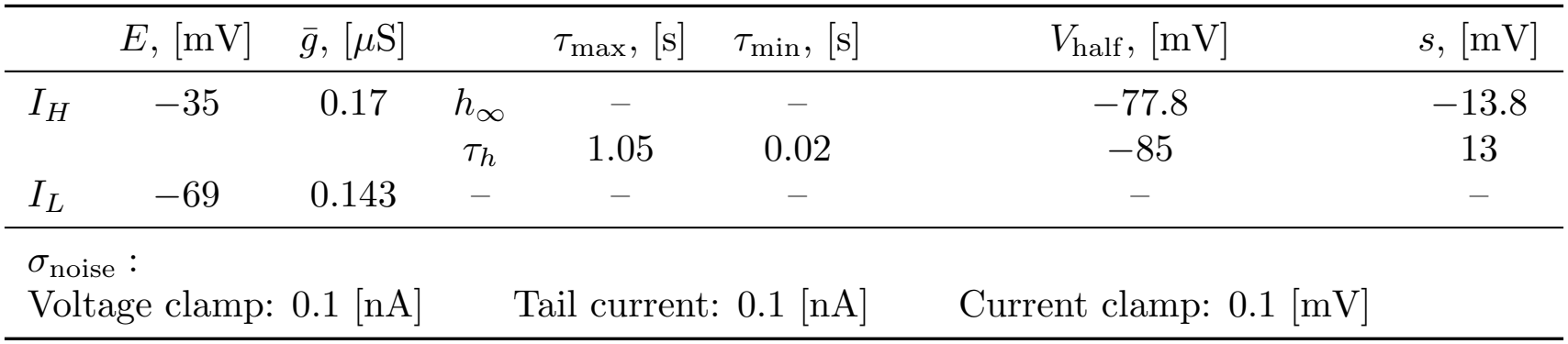
True parameters for the LGMD H-current model.

|  | *E*, [mV] | *ḡ*, [µS] |  | *τ*<sub>max</sub>, [s] | *τ*<sub>min</sub>, [s] | *V*<sub>half</sub>, [mV] | *s*, [mV] |
|---|---|---|---|---|---|---|---|
| *I*<sub>*H*</sub> | −35 | 0.17 | *h*<sub>∞</sub> | – | – | −77.8 | −13.8 |
|  |  |  | *τ*<sub>*h*</sub> | 1.05 | 0.02 | −85 | 13 |
| *I*<sub>*L*</sub> | −69 | 0.143 | – | – | – | – | – |

*σ*<sub>noise</sub> :
Voltage clamp: 0.1 [nA] Tail current: 0.1 [nA] Current clamp: 0.1 [mV]

**Table S2.**
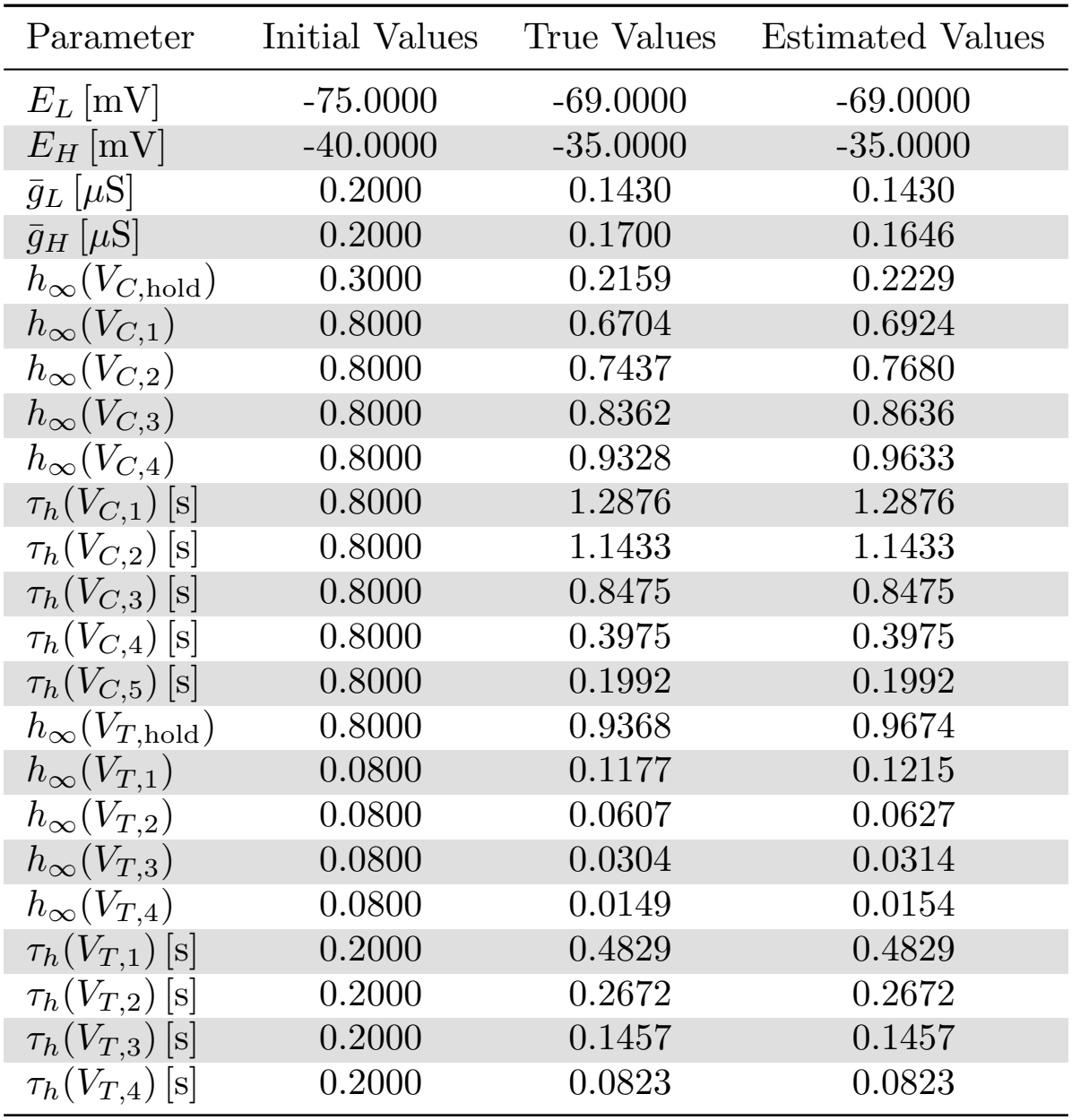
Parameter estimation results for the LGMD H-current model using noise-free voltage clamp, tail current, and current clamp data.

| Parameter | Initial Values | True Values | Estimated Values |
|---|---|---|---|
| E<sub>L</sub> [mV] | -75.0000 | -69.0000 | -69.0000 |
| E<sub>H</sub> [mV] | -40.0000 | -35.0000 | -35.0000 |
| ḡ<sub>L</sub> [µS] | 0.2000 | 0.1430 | 0.1430 |
| ḡ<sub>H</sub> [µS] | 0.2000 | 0.1700 | 0.1646 |
| h<sub>∞</sub>(V<sub>C,hold</sub>) | 0.3000 | 0.2159 | 0.2229 |
| h<sub>∞</sub>(V<sub>C,1</sub>) | 0.8000 | 0.6704 | 0.6924 |
| h<sub>∞</sub>(V<sub>C,2</sub>) | 0.8000 | 0.7437 | 0.7680 |
| h<sub>∞</sub>(V<sub>C,3</sub>) | 0.8000 | 0.8362 | 0.8636 |
| h<sub>∞</sub>(V<sub>C,4</sub>) | 0.8000 | 0.9328 | 0.9633 |
| τ<sub>h</sub>(V<sub>C,1</sub>) [s] | 0.8000 | 1.2876 | 1.2876 |
| τ<sub>h</sub>(V<sub>C,2</sub>) [s] | 0.8000 | 1.1433 | 1.1433 |
| τ<sub>h</sub>(V<sub>C,3</sub>) [s] | 0.8000 | 0.8475 | 0.8475 |
| τ<sub>h</sub>(V<sub>C,4</sub>) [s] | 0.8000 | 0.3975 | 0.3975 |
| τ<sub>h</sub>(V<sub>C,5</sub>) [s] | 0.8000 | 0.1992 | 0.1992 |
| h<sub>∞</sub>(V<sub>T,hold</sub>) | 0.8000 | 0.9368 | 0.9674 |
| h<sub>∞</sub>(V<sub>T,1</sub>) | 0.0800 | 0.1177 | 0.1215 |
| h<sub>∞</sub>(V<sub>T,2</sub>) | 0.0800 | 0.0607 | 0.0627 |
| h<sub>∞</sub>(V<sub>T,3</sub>) | 0.0800 | 0.0304 | 0.0314 |
| h<sub>∞</sub>(V<sub>T,4</sub>) | 0.0800 | 0.0149 | 0.0154 |
| τ<sub>h</sub>(V<sub>T,1</sub>) [s] | 0.2000 | 0.4829 | 0.4829 |
| τ<sub>h</sub>(V<sub>T,2</sub>) [s] | 0.2000 | 0.2672 | 0.2672 |
| τ<sub>h</sub>(V<sub>T,3</sub>) [s] | 0.2000 | 0.1457 | 0.1457 |
| τ<sub>h</sub>(V<sub>T,4</sub>) [s] | 0.2000 | 0.0823 | 0.0823 |

**Table S3.**
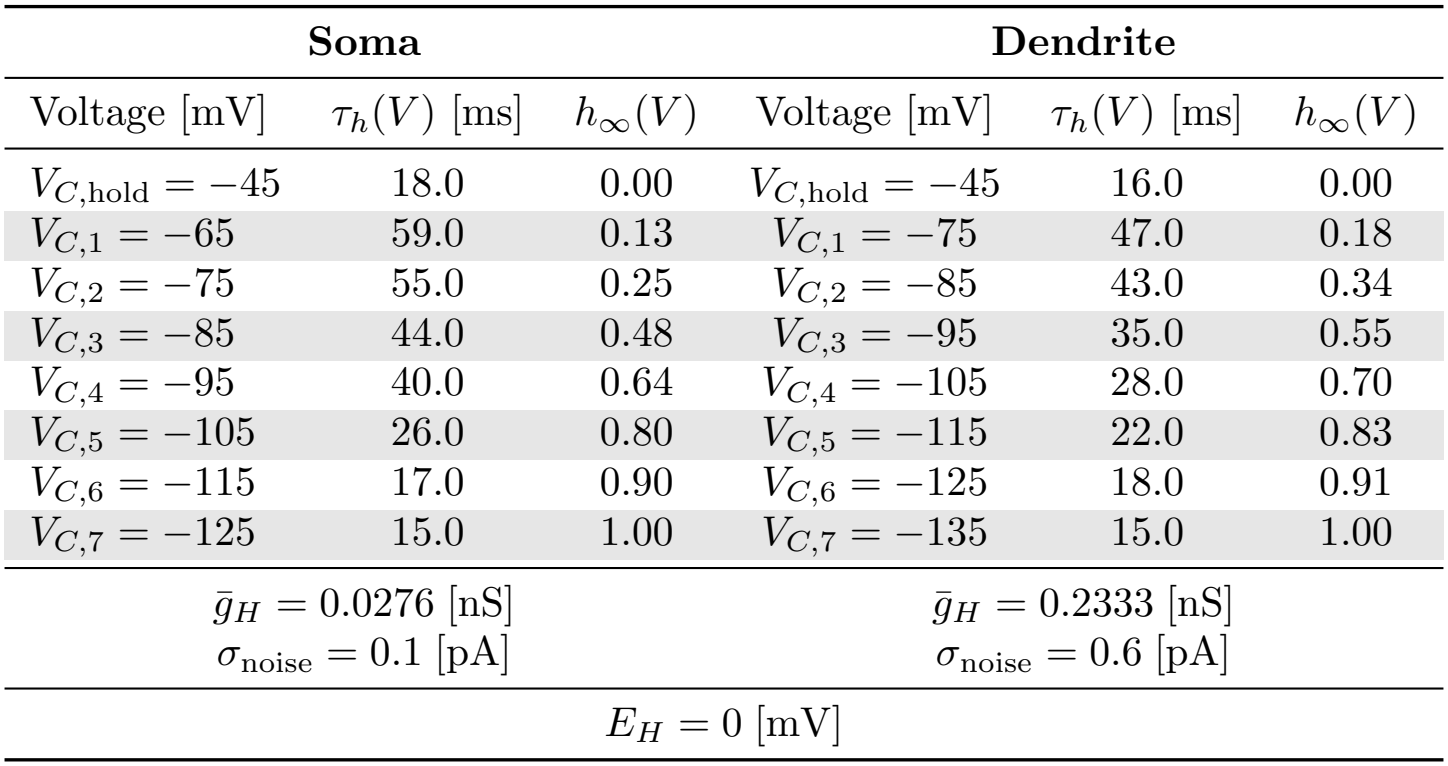
True parameters for the CA1 H-current model.

| Soma |  |  | Dendrite |  |  |
|---|---|---|---|---|---|
| Voltage [mV] | τ<sub>h</sub>(V) [ms] | h<sub>∞</sub>(V) | Voltage [mV] | τ<sub>h</sub>(V) [ms] | h<sub>∞</sub>(V) |
| V<sub>C,hold</sub> = −45 | 18.0 | 0.00 | V<sub>C,hold</sub> = −45 | 16.0 | 0.00 |
| V<sub>C,1</sub> = −65 | 59.0 | 0.13 | V<sub>C,1</sub> = −75 | 47.0 | 0.18 |
| V<sub>C,2</sub> = −75 | 55.0 | 0.25 | V<sub>C,2</sub> = −85 | 43.0 | 0.34 |
| V<sub>C,3</sub> = −85 | 44.0 | 0.48 | V<sub>C,3</sub> = −95 | 35.0 | 0.55 |
| V<sub>C,4</sub> = −95 | 40.0 | 0.64 | V<sub>C,4</sub> = −105 | 28.0 | 0.70 |
| V<sub>C,5</sub> = −105 | 26.0 | 0.80 | V<sub>C,5</sub> = −115 | 22.0 | 0.83 |
| V<sub>C,6</sub> = −115 | 17.0 | 0.90 | V<sub>C,6</sub> = −125 | 18.0 | 0.91 |
| V<sub>C,7</sub> = −125 | 15.0 | 1.00 | V<sub>C,7</sub> = −135 | 15.0 | 1.00 |
| ḡ<sub>H</sub> = 0.0276 [nS]<br>σ<sub>noise</sub> = 0.1 [pA] |  |  | ḡ<sub>H</sub> = 0.2333 [nS]<br>σ<sub>noise</sub> = 0.6 [pA] |  |  |
| E<sub>H</sub> = 0 [mV] |  |  |  |  |  |

**Table S4.**
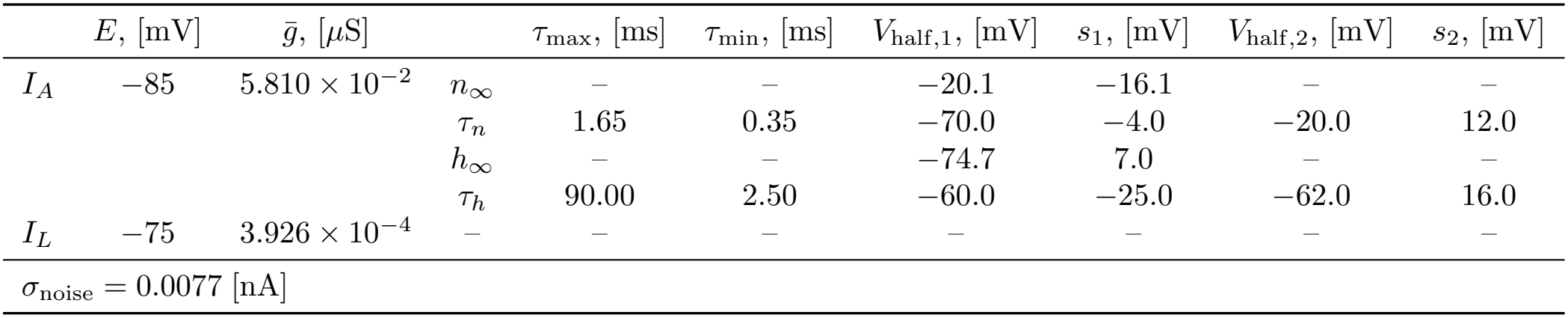
True parameters for the A-type current.

|  | E, [mV] | ḡ, [μS] |  | τ<sub>max</sub>, [ms] | τ<sub>min</sub>, [ms] | V<sub>half,1</sub>, [mV] | s<sub>1</sub>, [mV] | V<sub>half,2</sub>, [mV] | s<sub>2</sub>, [mV] |
|---|---|---|---|---|---|---|---|---|---|
| I<sub>A</sub> | −85 | 5.810 × 10<sup>−2</sup> | n<sub>∞</sub> | – | – | −20.1 | −16.1 | – | – |
|  |  |  | τ<sub>n</sub> | 1.65 | 0.35 | −70.0 | −4.0 | −20.0 | 12.0 |
|  |  |  | h<sub>∞</sub> | – | – | −74.7 | 7.0 | – | – |
|  |  |  | τ<sub>h</sub> | 90.00 | 2.50 | −60.0 | −25.0 | −62.0 | 16.0 |
| I<sub>L</sub> | −75 | 3.926 × 10<sup>−4</sup> | – | – | – | – | – | – | – |
| σ<sub>noise</sub> = 0.0077 [nA] |  |  |  |  |  |  |  |  |  |

**Table S5.** True parameters for the full Hodgkin-Huxley model.

|  | *E*, [mV] | *ḡ*, [mS/cm<sup>2</sup>] |
|---|---|---|
| *I*<sub>*Na*</sub> | 50.00 | 120.0 |
| *I*<sub>*K*</sub> | −77.00 | 36.0 |
| *I*<sub>*L*</sub> | −54.38 | 0.3 |

## Acknowledgments

We thank Dr. Jeff Magee for helpful discussions. We also thank Drs. Paul Pfaffinger and Richard Dewell for a careful reading of the manuscript.

Claude (Anthropic) and ChatGPT (OpenAI) were used for limited assistance in reviewing the manuscript, polishing the language of selected passages, and checking a few Jacobian rank calculations. All outputs were verified by the authors, who take full responsibility for the content of this article.

## References

1. Hodgkin AL, Huxley AF. A quantitative description of membrane current and its application to conduction and excitation in nerve. The Journal of Physiology. 1952;117(4):500–44. doi:10.1113/jphysiol.1952.sp004764.

2. Tononi G, Sporns O, Edelman GM. Measures of degeneracy and redundancy in biological networks. PNAS. 1999;96(6):3257–62. doi:10.1073/pnas.96.6.3257.

3. Albantakis L, Bernard C, Brenner N, Marder E, Narayanan R. The Brain’s Best Kept Secret Is Its Degenerate Structure. Journal of Neuroscience. 2024;44(40). doi:10.1523/JNEUROSCI.1339-24.2024.

4. Golowasch J, Abbott LF, Marder E. Activity-dependent regulation of potassium currents in an identified neuron of the stomatogastric ganglion of the crab Cancer borealis. Journal of Neuroscience. 1999;19(20):RC33. doi:10.1523/JNEUROSCI.19-20-j0004.1999.

5. Schulz DJ, Goaillard JM, Marder E. Variable channel expression in identified single and electrically coupled neurons in different animals. Nature Neuroscience. 2006;9(3):356–62. doi:10.1038/nn1639.

6. Marder E, Goaillard JM. Variability, compensation and homeostasis in neuron and network function. Nature Reviews Neuroscience. 2006;7(7):563–74. doi:10.1038/nrn1949.

7. MacLean JN, Zhang Y, Johnson BR, Harris-Warrick RM. Activity-independent homeostasis in rhythmically active neurons. Neuron. 2003;37(1):109–20. doi:10.1016/s0896-6273(02)01104-2.

8. Schulz DJ, Goaillard JM, Marder E. Quantitative expression profiling of identified neurons reveals cell-specific constraints on highly variable levels of gene expression. PNAS. 2007;104(32):13187–91. doi:10.1073/pnas.0705827104.

9. Goldman MS, Golowasch J, Marder E, Abbott LF. Global structure, robustness, and modulation of neuronal models. Journal of Neuroscience. 2001;21(14):5229–38. doi:10.1523/JNEUROSCI.21-14-05229.2001.

10. Calabrese RL, Prinz AA. Realistic Modeling of Small Neuronal Networks. In: De Schutter E, editor. Computational Modeling Methods for Neuroscientists. The MIT Press; 2009. p. 285–316. doi:10.7551/mitpress/7543.003.0014.

11. Prinz AA, Bucher D, Marder E. Similar network activity from disparate circuit parameters. Nature Neuroscience. 2004;7(12):1345–32. doi:10.1038/nn1352.

12. Brandoit J, Ernst D, Drion G, Fyon A. Fast reconstruction of degenerate populations of conductance-based neuron models from spike times. PLOS Comput Biol. 2026;22(5):e1014337. doi:10.1371/journal.pcbi.1014337.

13. Covarrubias M, Wei A, Salkoff L. Shaker, Shal, Shab, and Shaw express independent K+ current systems. Neuron. 1991 Nov;7(5):763–73. doi:10.1016/0896-6273(91)90279-9.

14. Bhalla T, Rosenthal JJC, Holmgren M, Reenan R. Control of human potassium channel inactivation by editing of a small mRNA hairpin. Nature Structural & Molecular Biology. 2004 Oct;11(10):950–6. doi:10.1038/nsmb825.

15. Schulte U, Thumfart JO, Klöcker N, Sailer CA, Bildl W, Biniossek M, et al. The Epilepsy-Linked Lgi1 Protein Assembles into Presynaptic Kv1 Channels and Inhibits Inactivation by Kvβ1. Neuron. 2006 Mar;49(5):697–706. doi:10.1016/j.neuron.2006.01.033.

16. Jindal HK, Folco EJ, Liu GX, Koren G. Posttranslational modification of voltage-dependent potassium channel Kv1.5: COOH-terminal palmitoylation modulates its biological properties. American Journal of Physiology-Heart and Circulatory Physiology. 2008 May;294(5):H2012–21. doi:10.1152/ajpheart.01374.2007.

17. Bernard C. Brain’s Best Kept Secret: Degeneracy. eNeuro. 2023 Nov;10(11). doi:10.1523/ENEURO.0430-23.2023.

18. Dewell RB, Gabbiani F. Biophysics of object segmentation in a collision-detecting neuron. eLife. 2018;7:e34238. doi:10.7554/eLife.34238.

19. Magee JC. Dendritic Hyperpolarization-Activated Currents Modify the Integrative Properties of Hippocampal CA1 Pyramidal Neurons. Journal of Neuroscience. 1998;18(19):7613–24. doi:10.1523/JNEUROSCI.18-19-07613.1998.

20. Wustenberg DG, Boytcheva M, Grunewald B, Byrne JH, Menzel R, Baxter DA. Current-and voltage-clamp recordings and computer simulations of Kenyon cells in the honeybee. Journal of Neurophysiology. 2004;92(4):2589–603. doi:10.1152/jn.01259.2003.

21. Heinkenschloss M, Namazifard S, Gabbiani F. Rank Deficiency in Parameter Estimation for Hodgkin-Huxley Type Current Models. Rice University: Department of Computational Applied Mathematics and Operations Research; 2026.

22. Golub GH, Van Loan CF. Matrix Computations. 4th ed. Johns Hopkins Studies in the Mathematical Sciences. Baltimore, MD: Johns Hopkins University Press; 2013. doi:10.56021/9781421407944.

23. Angelo K, London M, Christensen SR, Häusser M. Local and Global Effects of Ih Distribution in Dendrites of Mammalian Neurons. Journal of Neuroscience. 2007;27(32):8643–53. doi:10.1523/JNEUROSCI.5284-06.2007.

24. Pelz C, Jander J, Rosenboom H, Hammer M, Menzel R. IA in Kenyon Cells of the Mushroom Body of Honeybees Resembles Shaker Currents: Kinetics, Modulation by K+, and Simulation. Journal of Neurophysiology. 1999;81(4):1749–59. doi:10.1152/jn.1999.81.4.1749.

25. Hodgkin AL, Huxley AF. Currents carried by sodium and potassium ions through the membrane of the giant axon of Loligo. The Journal of Physiology. 1952;116(4):449–72. doi:10.1113/jphysiol.1952.sp004717.

26. Hansen PC. Rank-Deficient and Discrete Ill-Posed Problems. Numerical Aspects of Linear Inversion. SIAM Monographs on Mathematical Modeling and Computation. Society for Industrial and Applied Mathematics (SIAM), Philadelphia, PA; 1998. doi:10.1137/1.9780898719697.

27. Björck Å. Numerical Methods for Least Squares Problems. 2nd ed. Society for Industrial and Applied Mathematics (SIAM), Philadelphia, PA; 2024. doi:10.1137/1.9781611977950.

28. Straznicky JI, Gibson LA, Celse B, McAuley KB. Bayesian and subset-selection methods for parameter estimation in mechanistic models with limited data: A review and comparison. Chemical Engineering Research and Design. 2025;216:293–311. doi:10.1016/j.cherd.2025.02.037.

29. Vavoulis DV, Straub VA, Aston JAD, Feng J. A self-organizing state-space-model approach for parameter estimation in Hodgkin-Huxley-type models of single neurons. PLoS Comput Biol. 2012;8(3):e1002401. doi:10.1371/journal.pcbi.1002401.

30. Allgower EL, Georg K. Introduction to Numerical Continuation Methods. vol. 45 of Classics in Applied Mathematics. Society for Industrial and Applied Mathematics (SIAM), Philadelphia, PA; 2003. Reprint of the 1990 edition [Springer-Verlag, Berlin]. doi:10.1137/1.9780898719154.

31. Beyn WJ, Champneys A, Doedel E, Govaerts W, Kuznetsov YA, Sandstede B. Numerical continuation, and computation of normal forms. In: Handbook of dynamical systems, Vol. 2. North-Holland, Amsterdam; 2002. p. 149–219. doi:10.1016/S1874-575X(02)80025-X.

32. Doedel EJ. Lecture notes on numerical analysis of nonlinear equations. In: Krauskopf B, Osinga HM, Galán-Vioque J, editors. Numerical Continuation Methods for Dynamical Systems. Path Following and Boundary Value Problems. Underst. Complex Syst. Springer, Dordrecht; 2007. p. 1–49. doi:10.1007/978-1-4020-6356-51.

33. Ori H, Marder E, Marom S. Cellular function given parametric variation in the Hodgkin and Huxley model of excitability. Proc Natl Acad Sci USA. 2018;115(35):E8211–8. doi:10.1073/pnas.1808552115.

34. Therien AG, Blostein R. Mechanisms of sodium pump regulation. American Journal of Physiology-Cell Physiology. 2000 Sep;279(3):C541–66. doi:10.1152/ajpcell.2000.279.3.C541.

35. Contreras SA, Schleimer JH, Gulledge AT, Schreiber S. Activity-mediated accumulation of potassium induces a switch in firing pattern and neuronal excitability type. PLOS Computational Biology. 2021 May;17(5):e1008510. doi:10.1371/journal.pcbi.1008510.

36. Pressey JC, de Saint-Rome M, Raveendran VA, Woodin MA. Chloride transporters controlling neuronal excitability. Physiological Reviews. 2023 Apr;103(2):1095–135. doi:10.1152/physrev.00025.2021.

37. Csercsik D, Hangos KM, Szederkényi G. Identifiability analysis and parameter estimation of a single Hodgkin-Huxley type voltage dependent ion channel under voltage step measurement conditions. Neurocomputing. 2012;77:178–88. doi:10.1016/j.neucom.2011.09.006.

38. Walch OJ, Eisenberg MC. Parameter identifiability and identifiable combinations in generalized Hodgkin-Huxley models. Neurocomputing. 2016;199:137–43. doi:10.1016/j.neucom.2016.03.027.

39. Willms AR, Baro DJ, Harris-Warrick RM, Guckenheimer J. An Improved Parameter Estimation Method for Hodgkin-Huxley Models. J Comput Neurosci. 1999;6:145–68. doi:10.1023/A:1008880518515.

40. Korngreen A. Neuronal excitability and parameter variability in the Hodgkin-Huxley model. PLOS Comput Biol. 2026;22(6):e1014458. doi:10.1371/journal.pcbi.1014458.

41. Olypher AV, Calabrese RL. Using Constraints on Neuronal Activity to Reveal Compensatory Changes in Neuronal Parameters. Journal of Neurophysiology. 2007;98(6):3749–58. doi:10.1152/jn.00842.2007.

42. Raue A, Kreutz C, Maiwald T, Bachmann J, Schilling M, Klingmüller U, et al. Structural and practical identifiability analysis of partially observed dynamical models by exploiting the profile likelihood. Bioinformatics. 2009 Aug;25(15):1923–9. doi:10.1093/bioinformatics/btp358.

43. Hille B. Ion Channels of Excitable Membranes. 3rd ed. Oxford, UK: Oxford University Press; 2001. Available from: https://global.oup.com/ukhe/product/ion-channels-of-excitable-membranes-9780197583074.

44. Biegler LT. Nonlinear programming strategies for dynamic chemical process optimization. Theor Found Chem Eng. 2014;48(5):541–54. doi:10.1134/S0040579514050157.

45. Bard Y. Nonlinear Parameter Estimation. Academic Press [Harcourt Brace Jovanovich, Publishers], New York-London; 1974.

46. Dennis, Jr JE, Schnabel RB. Numerical Methods for Unconstrained Optimization and Nonlinear Equations. vol. 16 of Classics in Applied Mathematics. Society for Industrial and Applied Mathematics (SIAM), Philadelphia, PA; 1996. Corrected reprint of the 1983 original. doi:10.1137/1.9781611971200.

47. Madsen K, Nielsen HB. Introduction to Optimization and Data Fitting. DTU Informatics - IMM, Technical University of Denmark; 2010. Available at https://www2.imm.dtu.dk/pubdb/edoc/imm5938.pdf (accessed July 22, 2026).

48. Nocedal J, Wright SJ. Numerical Optimization. 2nd ed. Berlin, Heidelberg, New York: Springer Verlag; 2006. doi:10.1007/978-0-387-40065-5.

49. Virtanen P, Gommers R, Oliphant TE, Haberland M, Reddy T, Cournapeau D, et al. SciPy 1.0: Fundamental Algorithms for Scientific Computing in Python. Nature Methods. 2020;17:261–72. doi:10.1038/s41592-019-0686-2.

50. Newville M, Otten R, Nelson A, Stensitzki T, Ingargiola A, Allan D, et al. LMFIT: Non-Linear Least-Squares Minimization and Curve-Fitting for Python (Version 1.3.4). Zenodo. 2025. doi:10.5281/zenodo.16175987.

51. Moré JJ, Garbow BS, Hillstrom KE. User Guide for MINPACK 1. Argonne, Ill.: Argonne National Laboratory; 1980. doi:10.2172/6997568.

52. Moré JJ. The Levenberg–Marquardt Algorithm: Implementation and Theory. In: Watson GA, editor. Numerical Analysis, Proceedings, Biennial Conference, Dundee 1977. Berlin, Heidelberg, New-York: Springer Verlag; 1978. p. 105–16. doi:10.1007/BFb0067700.

53. Branch MA, Coleman TF, Li Y. A subspace, interior, and conjugate gradient method for large-scale bound-constrained minimization problems. SIAM J Sci Comput. 1999;21(1). doi:10.1137/S1064827595289108.

54. The JAX Authors. JAX: High performance array computing (Version 0.9.2); 2026. Available from: https://docs.jax.dev/en/latest.

55. Lee JM. Introduction to Smooth Manifolds. vol. 218 of Graduate Texts in Mathematics. 2nd ed. Springer, New York; 2013. doi:10.1007/978-1-4419-9982-5.

56. Conlon L. Differentiable Manifolds. 2nd ed. Modern Birkhäuser Classics. Birkhäuser Boston, Inc., Boston, MA; 2008. doi:10.1007/978-0-8176-4767-4.

57. Donaldson JR, Schnabel RB. Computational experience with confidence regions and confidence intervals for nonlinear least squares. Technometrics. 1987;29(1):67–82. doi:10.2307/1269884.

58. Seber GAF, Wild CJ. Nonlinear Regression. Wiley-Interscience Paperback Series. John Wiley & Sons, Inc., New York; 2003. doi:10.1002/0471725315.

